# Determination of hydrogen bonds in Gromacs: new implementation to overcome the limitation

**DOI:** 10.1101/2023.09.01.555860

**Authors:** S.V. Gorelov, A. I. Titov, O.A. Tolicheva, A. L. Konevega, A. V. Shvetsov

## Abstract

This work describes a new software algorithm associated with determination of hydrogen bonds in biomacromolecules and their environment. The already existing algorithm for determining hydrogen bond networks in Gromacs has a number of impenetrable limitations in the analysis of structures and trajectories. The new implementation of the algorithm for determining hydrogen bond networks in the form of a native Gromacs trajectory analysis module allows to quickly analyze molecular dynamics trajectories without restrictions, thereby overcoming the limitation of original algorithm.

The application of the developed algorithm made it possible to obtain and analyze the networks of hydrogen bonds of the studied defensin-like protein Pentadiplandra brazzeana, as well as to study the lifetime of hydrogen bonds between the residues of this protein and water molecules. The data obtained using the new implementation coincided with the experimental one.

**TOC Graphic:** 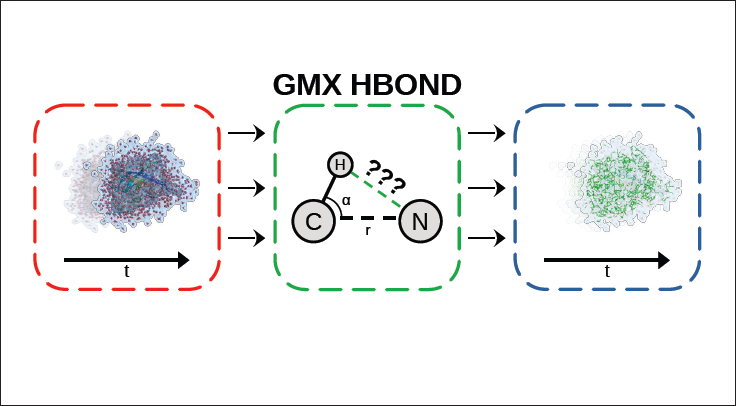

## Introduction

A hydrogen bond is a type of attractive (dipole-dipole) interaction between an electronegative atom and a hydrogen atom bonded to another electronegative atom. This bond always involves a hydrogen atom. Such bonds can occur between molecules or within parts of a single molecule.^1^ They provide most of the directional interactions that underlie protein folding, protein secondary structure, and molecular recognition, and thus are essential for understanding protein spatial organization and movement. The energy of such bonds is experimentally known, which varies greatly from *≈*5–6 kcal/mol for an isolated bond, to *≈*0.5–1.5 kcal/mol for proteins in solution. The cores of most protein structures consist of secondary structures such as the *α*-helices and the *β*-sheets. This satisfies the hydrogen binding potential between the carbonyl oxygen of the main chain and the nitrogen of the amino group embedded in the hydrophobic core of the protein. The hydrogen bond between a protein and its ligands (which can be biomacromolecules such as proteins, nucleic acids, substrates, effectors, or inhibitors) ensures the directionality and specificity of the interaction, which is a fundamental aspect of molecular recognition. Therefore, the energetics and kinetics of the hydrogen bond must be optimal to allow fast selection and folding kinetics, conferring stability on the protein structure and providing the specificity required for selective macromolecular interactions. ^2,3^

An implementation of the algorithm for determining hydrogen bond networks can be used as a Gromacs trajectory analysis module to unambiguously determine the existing hydrogen bonds in a trajectory. Knowledge of existing hydrogen bonds can be used, for example, to find protein residues that actively bind to water, to evaluate protein packaging and stabilization, to search for protein secondary structures, or to evaluate interactions with ligands or at protein-protein interfaces.

## Implementation

In the newly developed implementation of the hydrogen bond network determination algorithm, these bonds served as the basis for the analysis, with oxygen or nitrogen atoms serving as acceptors, and oxygen or nitrogen covalently bonded with one or more hydrogen atoms serving as donors. To determine networks of hydrogen bonds in trajectories, a geometric criterion for the existence of a hydrogen bond is used:

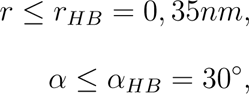

where *r* is the distance between the donor and the acceptor, and *α* is the Hydrogen-Donor-Acceptor angle. The value *r_HB_* = 0.35 nm corresponds to the first minimum of the radial distribution functions of water in the SPC/E model.^4^

**Figure 1:**
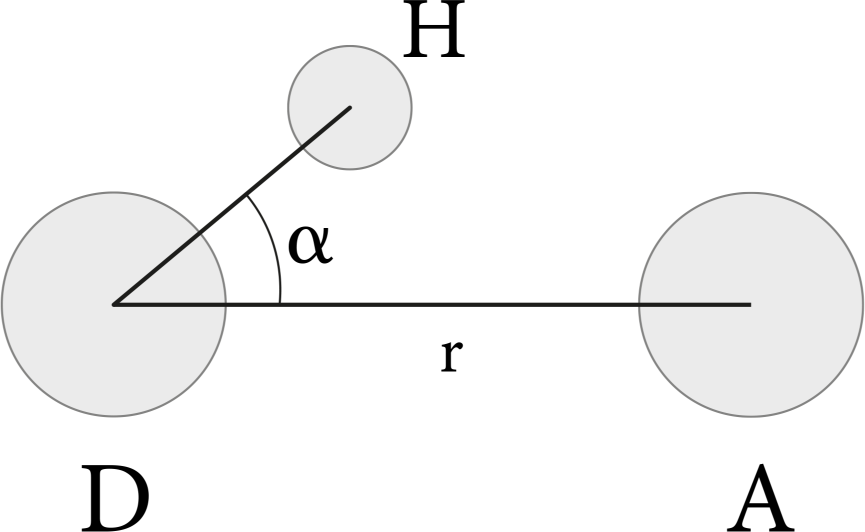
Geometric criterion for the existence of a hydrogen bond between donor D and acceptor A with the participation of the hydrogen atom H.

First of all, the algorithm analyzes the trajectory and topology received at the input, checking their correctness and compatibility with each other. Next, the algorithm asks the user to indicate two groups of atoms from the trajectory - reference and target - in order to analyze the existing hydrogen bonds between them. These two groups of atoms must be either absolutely identical or strictly non-intersecting. Otherwise, the algorithm will report an error and will not analyze the structure further.

If the topology of the trajectory and the selected groups have successfully passed all the checks, then the algorithm will begin to selection of potential donors and acceptors from topology based on the reference and target groups. All oxygen atoms are always considered as potential hydrogen bond acceptors, while nitrogen atoms can be considered as potential acceptors or ignored in the definition of acceptors (this depends on the parameter passed by the user “-an” or “-noan”; by default, nitrogen atoms are considered potential acceptors, corresponding to the choice of “-an”). Donors are defined in terms of interaction functions obtained from the topology. From the interaction functions of the SETTLE type, hydrogen donors for water molecules are determined, which can be exclusively only oxygen atoms. Of the other interaction functions that can determine the formation of a chemical bond strictly between two arbitrary atoms, oxygen and nitrogen atoms are selected. In addition, for each donor, the hydrogen atoms associated with it are memorized. After all donors and acceptors have been identified, there is a special designation of those acceptors that have also been identified as donors.

If no donor or acceptor was found in any of the groups — the algorithm will issue a warning. If no donor and no acceptor are found in any of the groups, the algorithm will terminate its work prematurely with a special message to the user, since from such a data set, where there are no critically important components (donors and acceptors), hydrogen bond analysis will not be possible.

Next, the existing hydrogen bonds are calculated according to the geometric criterion at each moment of the trajectory. A search for neighboring atoms is carried out using the Neighbor Search algorithm: the reference and target groups are divided into donors and acceptors, and then pairs of atoms are selected (acceptor atoms from the reference group are compared with donor atoms from the target group) located at a distance of no more than 0.35 nm (this value can also be set by the user, but it cannot be less than 0.35 nm). In these pairs, the geometric criterion is checked for the donor, acceptor, and all hydrogen atoms covalently bonded to the donor. If the criterion was satisfied, then the corresponding pairs with hydrogen are recorded in the storage. After the end of this procedure, the search starts again, but only if the reference group and the target group were not identical to each other. In the new round of the search, hydrogen bonds will be determined already between donors of the reference and acceptors of the target groups.

**Figure 2:**
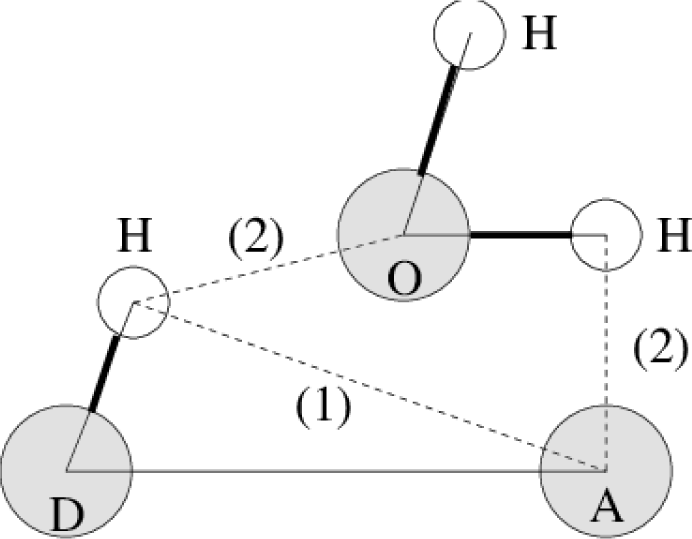
Hydrogen bond calculation. (1) Normal hydrogen bond between two residues. (2) A hydrogen bond bridge across a water molecule. ^4^

As a result (by default), a file will be provided with the indices of all atoms of the reference and target groups and the indices of all atoms that are donors and acceptors for these groups. It will also list all hydrogen bonds that exist in the trajectory. These bonds can (at the user’s request; parameter “-pf”) be specified for each frame separately, or they can (by default; “-nopf”) be specified for the entire structure as a whole (in this case, duplicated hydrogen links are not reflected in the output file). It is also possible (parameter “-m”) to combine information about hydrogen bonds, which, in the above approximation (for each frame, or for the entire trajectory), having identical donor and acceptor indices, will differ only in the hydrogen index. However, by default (parameter “-nom”) pairs of hydrogen bonds will not merge together and all found pairs will be presented in the output.

## Example applications

As a result, an algorithm for determining networks of hydrogen bonds was developed. The main output of the developed algorithm is an index map, which contains the indices of acceptors, donors, hydrogen atoms associated with donors, and all atoms that participate in the formation of a hydrogen bond. In addition to the index map, the algorithm is also capable of producing optional files that contain data for plot creation of certain key statistics.

The developed algorithm was applied to the molecular dynamics trajectory of the defensin-like protein of Pentadiplandra brazzeana (Brazzein). The duration of the trajectory is 100 ns.

## Discussion

From the time dependences of the number of hydrogen bond donors and acceptors (see Fig. 3), it becomes clear that in the trajectoriy of molecular dynamics in pairs protein-protein and, especially, in protein-water pairs, the number of hydrogen bond donors consistently exceeds the number of acceptors. As expected, in water-water pairs, on average, the number of donors is identical to the number of acceptors.

**Figure 3:**
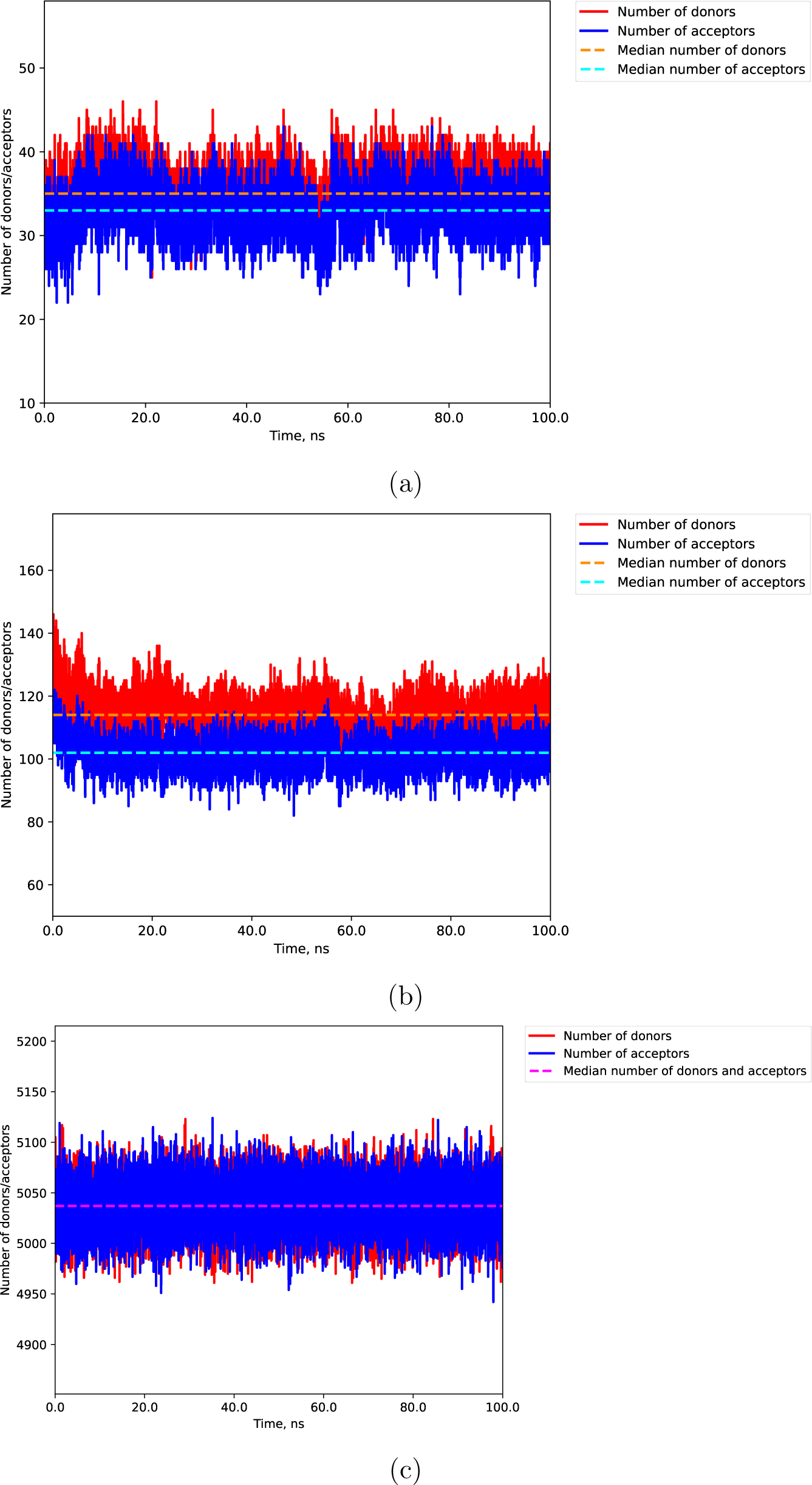
Graphs of the dependence of the number of donors and acceptors of a hydrogen bond between protein atoms (3a), between protein and water atoms (3b) and between water atoms (3c) as a function of time.

As can be seen from the obtained histograms of the distribution of hydrogen bond distances (see Fig. 6), the most probable values of hydrogen bond lengths are in the range 0 .24-0.35 nm. This is consistent with the theory of hydrogen bond formation, in which hydrogen bonds with a distance between the donor and acceptor of 0.22-0.25 nm are considered strong (mainly covalent), those with a distance of 0.25-0.32 nm are considered moderate (in mostly electrostatic), and those with distances exceeding 0.32 nm are considered weak (electrostatic).^5,6^ The presented graphs show that most of the identified hydrogen bonds in the studied trajectory are moderate. Weak and strong hydrogen bonds are also present in a smaller amount.

It is known^7,8^ that the distance between hydrogen and a hydrogen bond donor (if the donor is a nitrogen atom N or an oxygen atom O) is *≈*1 Å. It is also known that the closer the angle *∠_DHA_* to 180*^◦^*, the stronger the interatomic interactions inside the hydrogen bond (the bond is considered strong when the angle *∠_DHA_* reaches 170*^◦^*).^6^ From this we can conclude that the smaller the angle *∠_HDA_*, the stronger the hydrogen bond will be, and the interaction within the hydrogen bond will be especially strong at *∠_HDA_ ≤* 7.16*^◦^*. Weak hydrogen bonds can be considered as bonds with an angle *∠_HDA_* > 30*^◦^*, but, thus, they are eliminated when determining a hydrogen bond by a geometric criterion (see 1) and, therefore, are outside the range of values obtained by the software algorithm in the output file. Based on the obtained histograms of the distribution of hydrogen bond angles (see Fig. 5), one can see that hydrogen bonds of moderate strength are predominantly formed in all trajectoriy.

Using the developed algorithm, it was possible to determine the lifetime of hydrogen bonds between protein atoms and water atoms for each donor and acceptor in the studied trajectory (see Table 1). The obtained data was compared with the hydrogen bonds found in the crystal structures of the studied proteins obtained experimentally (using the X-ray diffraction method).

**Table 1:** List of residues involved in the formation of hydrogen bonds as donors of the defensin-like protein of Pentadiplandra brazzeana (Brazzein). Residues that form hydrogen bonds as donors in the crystal structure with identifier 7w8e are highlighted in bold.

| Hydrogen bond donor residues | Life time, ps |
| --- | --- |
| **K3** | 99260 |
| K15 | 98840 |
| **K6** | 98770 |
| **K27** | 98300 |
| D2 | 97750 |
| **K30** | 97300 |
| **K5** | 97100 |
| R43 | 96850 |
| R43 | 95600 |
| **R33** | 93590 |
| N20 | 93440 |
| S14 | 93250 |
| **Q17** | 92680 |
| K42 | 92590 |
| **Y39** | 91270 |
| **R33** | 90140 |
| **Y24** | 88900 |
| **N23** | 87750 |
| **C22** | 86040 |
| Y54 | 83430 |
| **Q21** | 82910 |
| R43 | 79400 |
| **Y39** | 76640 |
| **V7** | 75780 |
| **N10** | 74400 |
| A19 | 72250 |
| **K3** | 70610 |
| **H31** | 69870 |
| V13 | 67870 |
| N20 | 66650 |
| S14 | 65960 |

**Table 2:**
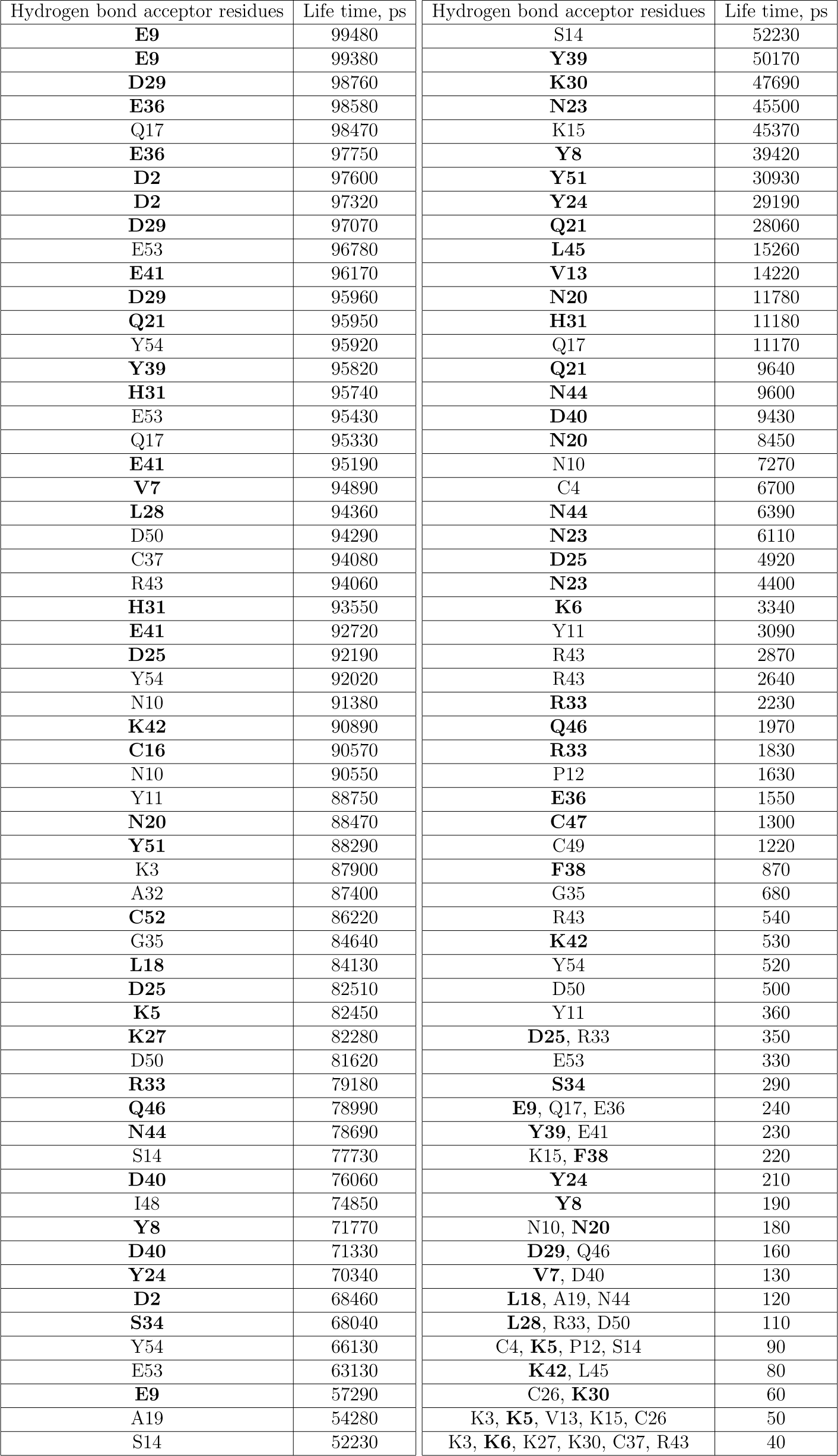
List of residues involved in the formation of hydrogen bonds as acceptors of the defensin-like protein of Pentadiplandra brazzeana (Brazzein). Residues that form hydrogen bonds as acceptors in the crystal structure with identifier 7w8e are highlighted in bold.

| Hydrogen bond acceptor residues | Life time, ps | Hydrogen bond acceptor residues | Life time, ps |
| --- | --- | --- | --- |
| **E9** | 99480 | S14 | 52230 |
| **E9** | 99380 | **Y39** | 50170 |
| **D29** | 98760 | **K30** | 47690 |
| **E36** | 98580 | **N23** | 45500 |
| Q17 | 98470 | K15 | 45370 |
| **E36** | 97750 | **Y8** | 39420 |
| **D2** | 97600 | **Y51** | 30930 |
| **D2** | 97320 | **Y24** | 29190 |
| **D29** | 97070 | **Q21** | 28060 |
| E53 | 96780 | **L45** | 15260 |
| **E41** | 96170 | **V13** | 14220 |
| **D29** | 95960 | **N20** | 11780 |
| **Q21** | 95950 | **H31** | 11180 |
| Y54 | 95920 | Q17 | 11170 |
| **Y39** | 95820 | **Q21** | 9640 |
| **H31** | 95740 | **N44** | 9600 |
| E53 | 95430 | **D40** | 9430 |
| Q17 | 95330 | **N20** | 8450 |
| **E41** | 95190 | N10 | 7270 |
| **V7** | 94890 | C4 | 6700 |
| **L28** | 94360 | **N44** | 6390 |
| D50 | 94290 | **N23** | 6110 |
| C37 | 94080 | **D25** | 4920 |
| R43 | 94060 | **N23** | 4400 |
| **H31** | 93550 | **K6** | 3340 |
| **E41** | 92720 | Y11 | 3090 |
| **D25** | 92190 | R43 | 2870 |
| Y54 | 92020 | R43 | 2640 |
| N10 | 91380 | **R33** | 2230 |
| **K42** | 90890 | **Q46** | 1970 |
| **C16** | 90570 | **R33** | 1830 |
| N10 | 90550 | P12 | 1630 |
| Y11 | 88750 | **E36** | 1550 |
| **N20** | 88470 | **C47** | 1300 |
| **Y51** | 88290 | C49 | 1220 |
| K3 | 87900 | **F38** | 870 |
| A32 | 87400 | G35 | 680 |
| **C52** | 86220 | R43 | 540 |
| G35 | 84640 | **K42** | 530 |
| **L18** | 84130 | Y54 | 520 |
| **D25** | 82510 | D50 | 500 |
| **K5** | 82450 | Y11 | 360 |
| **K27** | 82280 | **D25, R33** | 350 |
| D50 | 81620 | E53 | 330 |
| **R33** | 79180 | **S34** | 290 |
| **Q46** | 78990 | **E9, Q17, E36** | 240 |
| **N44** | 78690 | **Y39, E41** | 230 |
| S14 | 77730 | K15, **F38** | 220 |
| **D40** | 76060 | **Y24** | 210 |
| I48 | 74850 | **Y8** | 190 |
| **Y8** | 71770 | N10, **N20** | 180 |
| **D40** | 71330 | **D29, Q46** | 160 |
| **Y24** | 70340 | **V7, D40** | 130 |
| **D2** | 68460 | **L18, A19, N44** | 120 |
| **S34** | 68040 | **L28, R33, D50** | 110 |
| Y54 | 66130 | C4, **K5, P12, S14** | 90 |
| E53 | 63130 | **K42, L45** | 80 |
| **E9** | 57290 | C26, **K30** | 60 |
| A19 | 54280 | K3, **K5, V13, K15, C26** | 50 |
| S14 | 52230 | K3, **K6, K27, K30, C37, R43** | 40 |

It should be noted that some residues in the tables may be repeated, since they can contain several atoms that act as a hydrogen bond donor or acceptor. Therefore, each of these atoms can independently form hydrogen bonds. And since the lifetime of a hydrogen bond was determined specifically for each protein atom individually, the values of the formed hydrogen bonds for them were recorded individually.

All the residues (except for Ala43 residue — it was replaced by Arg43 in the Brazzein protein, examined in the trajectory) that were identified as containing atoms of donors and acceptors of hydrogen bonds from protein crystals also formed hydrogen bonds of varying degrees of stability (mostly stable, i.e. forming hydrogen bonds that exist for more than 2% (2 ns) of the time of the total duration (100 ns) of the trajectory).

From the Brazzein protein trajectory, it was possible to isolate many residues that had hydrogen bonding atoms in the crystal structure and yet also formed a relatively stable hydrogen bond with water through their atoms in a molecular dynamics trajectory. These are the residues: Asp2, Lys3, Cys4, Lys5, Lys6, Val7, Tyr8, Glu9, Asn10, Val13, Cys16, Gln17, Leu18, Asn20, Gln21, Cys22, Asn23, Tyr24, Asp25, Leu28, Asp29, Lys30, His31, Arg33, Ser34, Glu36, Phe38, Tyr39, Asp40, Glu41, Lys42, Asn44, Leu45, Gln46, Cys47, Tyr51, Cys52.

The residues with the longest-lived hydrogen bonds between protein and water were analyzed for the formation of stationary bonds. Of the most stable hydrogen bonds formed by residues in the trajectoriy of molecular dynamics, we selected those that formed a longterm bond (both as a donor and as an acceptor) with the same water molecule.

Table 3 shows that the Cys22 and Leu18 residues, which were previously identified as forming hydrogen bonds with water in the crystal structure with identifier 7w8e, in addition to forming stable hydrogen bonds in the molecular dynamics trajectory, also form long-lived stationary bonds with water molecules.

**Table 3:** List of Brazzein protein residues that forming stable stationary hydrogen bonds with water molecules. Residues forming hydrogen bonds in the crystal structure with identifier 7w8e are highlighted in bold.

| Residues with stationary connections | Longest lifetime, ps | Numer of stationary connections |
| --- | --- | --- |
| **C22** | 12870 | 11 |
| **L18** | 12170 | 12 |
| K15 | 7220 | 7 |
| **N23** | 4940 | 3 |
| G35 | 4860 | 3 |
| **E36** | 2160 | 1 |
| D50 | 2140 | 1 |
| **Q21** | 2010 | 1 |

Thus, using the developed algorithm for determining hydrogen bond networks, a conclusion was made not only regarding the general abilities to form hydrogen bonds in individual protein residues, but also regarding the ability to form long-lived stationary bonds with water molecules. The results obtained using the developed algorithm agree with the results obtained experimentally.

## Comparison of algorithms’ “run time”

The speed of the old and new implementations of hydrogen bond determination algorithms was tested on arbitrary trajectories of different sizes for defensin-like protein of Pentadiplandra brazzeana, Sars-Cov-2 Non-Structural Protein 12, T7 RNA polymerase and scFv antibodies: 3B12, 3H10, B11-1, B11-2 and ST4. In all cases, only protein and/or water atoms from the molecular dynamics trajectory were analyzed to obtain records of hydrogen bonds.

## Conclusions

The re-implementation of the hydrogen bond network determination algorithm in the trajectory of defensin-like protein of Pentadiplandra brazzeana molecular dynamics made it possible to obtain a detailed list of residues that are most actively involved in the formation of hydrogen bonds both as a donor (see Table 1) and as an acceptor (see Table 2) and determine that in the Brazzin protein residues Cys22 and Leu18 are the most active residues that form stationary hydrogen bonds.

Unlike the original algorithm for determining hydrogen bonds nerworks, the new implementation consumes much less memory and works much faster. The data obtained as a result of the work of the new algorithm do not differ from those obtained by the original algorithm. The new implementation is capable of accepting structures of any size as input, without restrictions, and can output information both about the contained hydrogen bonds at each moment of the trajectory, and about hydrogen bonds in the trajectory as a whole. In the new implementation, the shortcomings contained in the previous implementation of this algorithm were eliminated.

## Data Availability

Our standalone implementation of the algorithm for determining hydrogen bond networks in Gromacs is available at the GitLab repository: https://gitlab.com/bio-pnpi/gmx-hbond. You can also look at the measured numerical results of speed comparisons (in the form of graphs and tables) at https://gitlab.com/bio-pnpi/gmx-hbond-refdata.

## Acknowledgments

The enhanced algorithm for determining networks of hydrogen bonds was developed and implemented within the state assignment of Ministry of Science and Higher Education of the Russian Federation (theme N*^o^* 121060200127-6 and theme N*^o^* 2.8629.2017). The structures and molecular dynamics trajectories of scFv antibodies were obtained within the state assignment of Ministry of Science and Higher Education of the Russian Federation (theme N*^o^* 075-15-2021-1360).

## Supporting Information Available

### Initial structures

Initial structures for test objects were taken from the following sources:

- structure of the defensin-like protein of *Pentadiplandra brazzeana* was taken from PDB ID 1BRZ
- structure of NSP12 from SARS-Cov-2 was taken from PDB ID 7BTF
- structure of T7 polymerase was taken from PDB ID 2PI5

Test objects 3B12, 3H10, B11-1, B11-2 and ST4 are scFv antibody. Seq sequences used to create them were constructed in a following way. Briefly, mice were immunized with recombinant VEGFR-1, CTLA-4 and Stabilin-1 proteins. Obtained B-lymphocytes were fused to myeloma cells and producing monoclonal antibodies hybridomas: 3B12 and 3H10 variants against VEGFR-1, B11-1 and B11-2 against CTLA-4 and St4 against Stabilin-1, were selected. The genes of the variable parts of heavy and light chains (VH and VL respectively) were sequenced and converted into amino acid sequences. Next, VH and VL fragments were connected into a single polypeptide chain via a (G4S)3 flexible linker in a «head-to-tail» manner, and tagged with a GSS-His6 at the C terminus. Thus, the final domain structure of these scFv antibodies is VH-(G4S)3-VL-GSS-His6. Each scFv antibody structure was modelled using AlphaFold.^9^

### Molecular dynamics

Molecular dynamics (MD) modeling was performed with the software package GROMACS 2023,^10^ the amber14sb field^11^ was used for the protein and tip3p was used as a water model. The resulting systems were placed in a periodic water box in such a way that at least 25Å remained to the walls of the box. Then the resulting box was minimized using the steepest descent algorithm. The system obtained as a result of minimization was subjected to charge neutralization by adding 150 mM of NaCl bringing the total charge of the system to zero. The neutralized system was again subjected to the procedure of energy minimization using the steepest descent algorithm. Next, the system was equilibrated using a two-stage approach. At the first stage, all heavy atoms of protein were restrained to their initial positions using an additional energy term (posres), while at the start of equilibration, the temperature (particle velocity distribution) was taken from the Maxwell distribution for a given temperature at 310K. The system was equilibrated for 5 ns at each temperature. The integration step was 2 fs; the V-rescale thermostat and the C-rescale barostat were used. At the second stage, the additional restraining potential was removed, and all components of the system could move freely. During this stage, the Nose-Hoover thermostat and the Parrinello-Rahman barostat were used, and the system was equilibrated for 10 ns. The final state obtained as a result of a two-stage equilibration was used as a start for the production dynamics. Molecular dynamics was carried out for 1000 ns, using the same set of parameters as for the second stage of equilibration.

### Example applications graphs

Below are the graphs mentioned in the “example applications” section. They are images of graphs based on data obtained using the developed algorithm for determining networks of hydrogen bonds.

**Figure 4:**
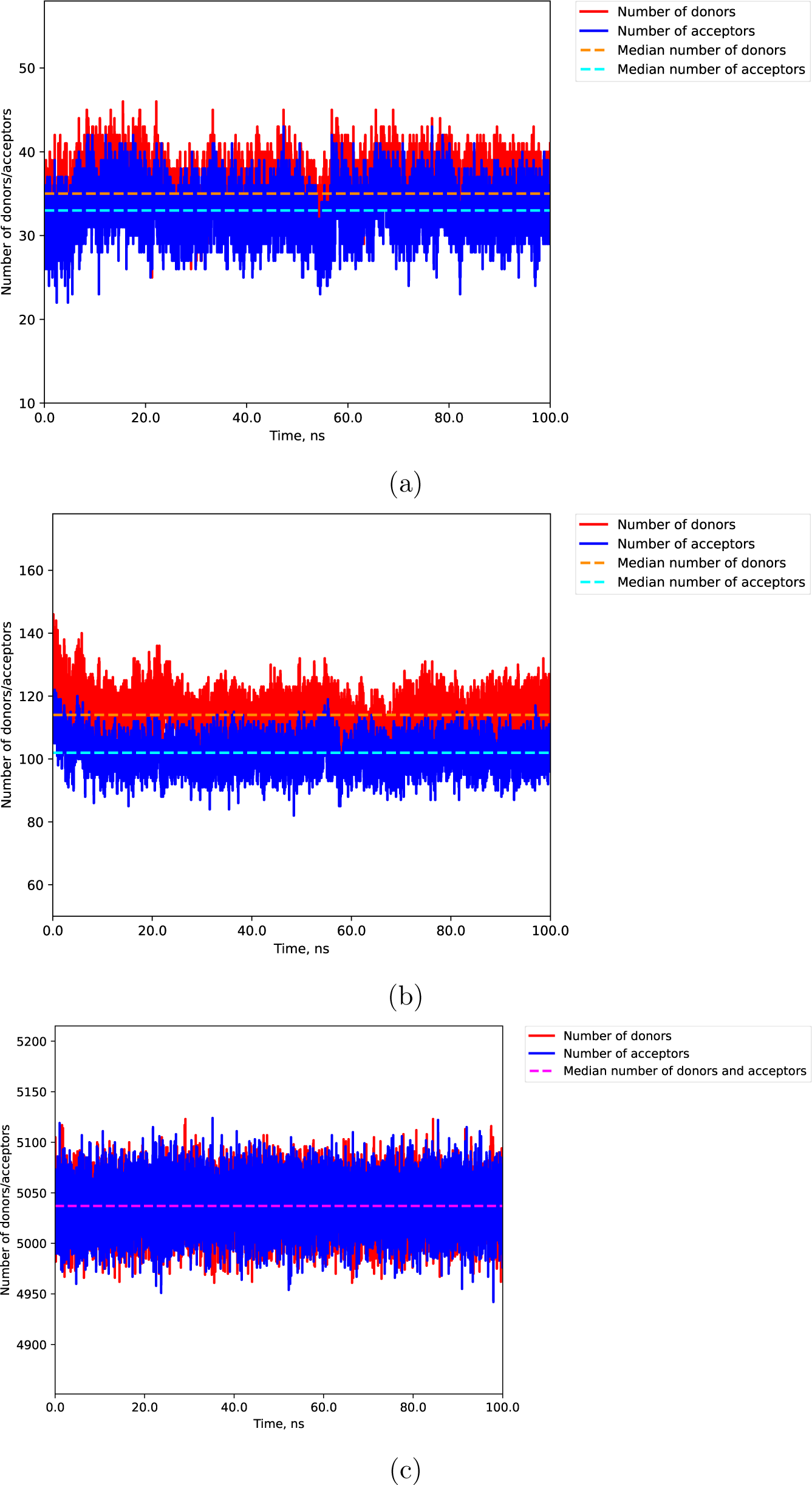
Graphs of the number of hydrogen bonds between protein atoms (4a), between protein and water atoms (4b) and between water atoms (4c) as a function of time.

**Figure 5:**
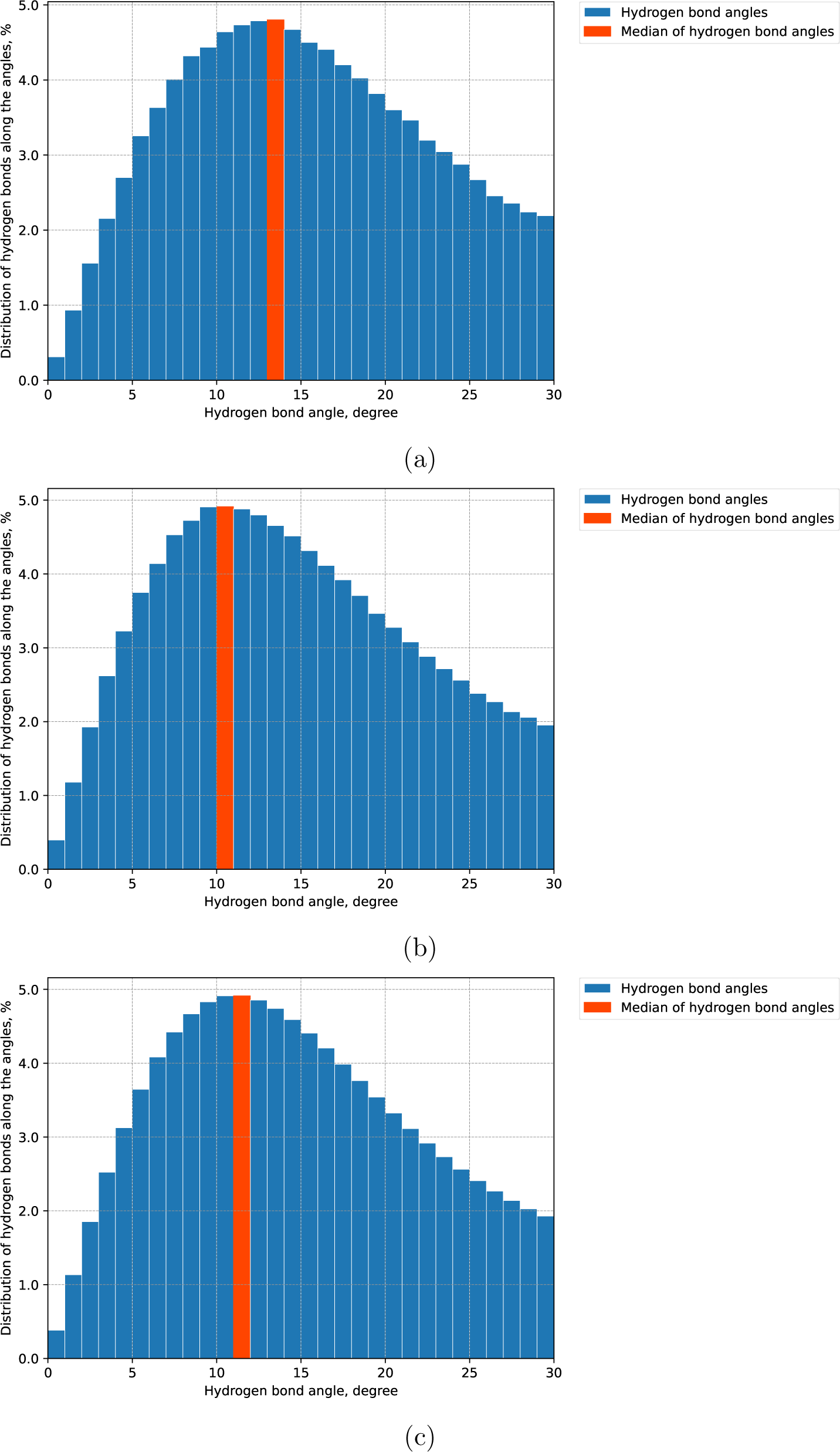
Histograms of distribution of angles of hydrogen bonds between protein atoms (5a), between protein and water atoms (5b) and between water atoms (5c).

**Figure 6:**
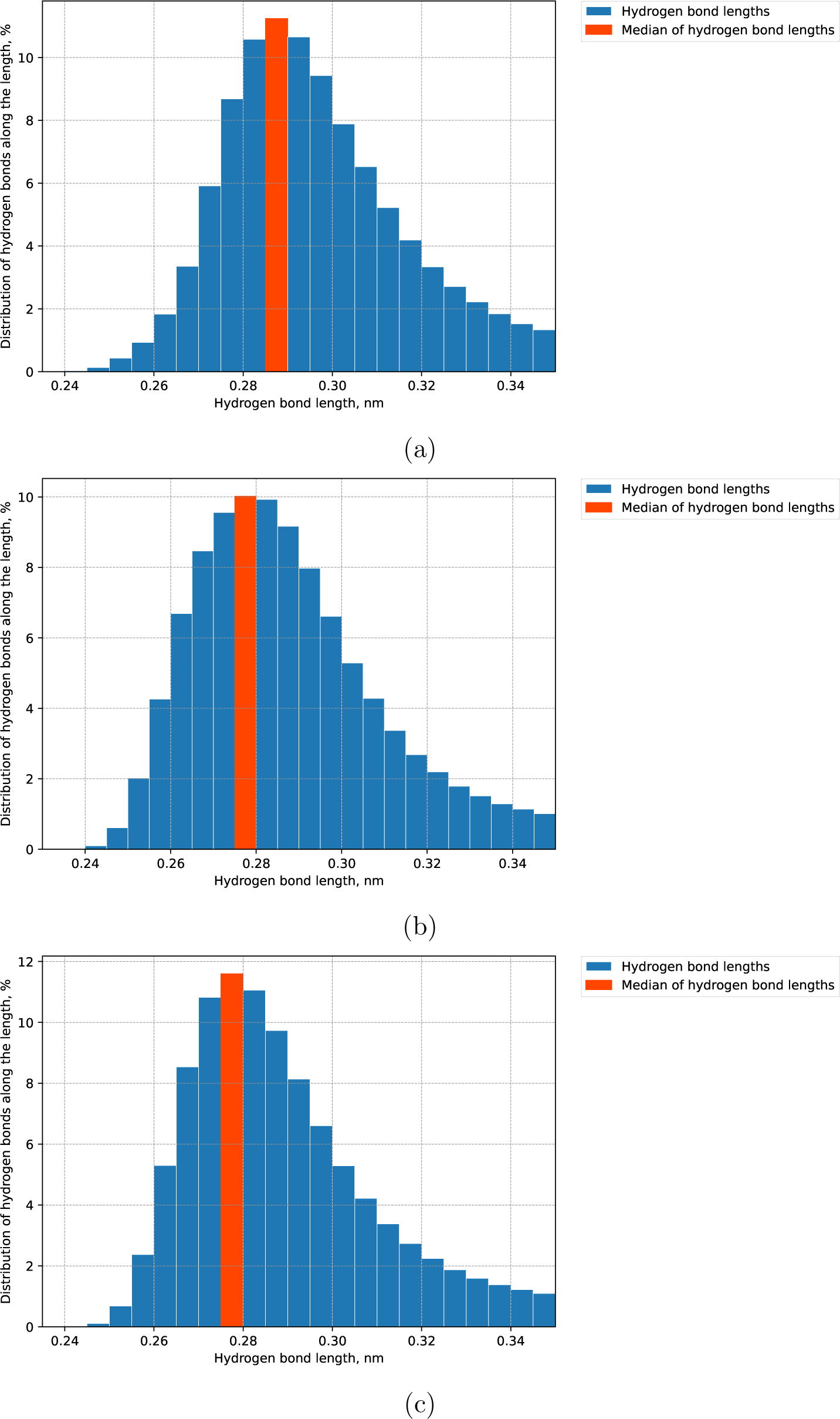
Histograms of the distribution of hydrogen bond distances between protein atoms (6a), between protein and water atoms (6b) and between water atoms (6c).

### Comparison of algorithms’ “run time”

The speeds of the old and new implementations of the algorithm for determining hydrogen bonds were tested on arbitrary trajectories of different sizes for the defensin-like protein of Pentadiplandra brazzeana, Sars-Cov-2 Non-Structural Protein 12, T7 RNA polymerase and scFv antibodies: 3B12, 3H10, B11-1, B11-2 and ST4. In all cases, the old implementation ran in one and 40 threads, while the new implementation ran in one thread. When testing the running time, each algorithm was given a trajectory with instructions to determine hydrogen bonds inside protein molecules, inside water molecules, and between protein and water molecules. You can also view the resulting graphs for all trajectories in the GitLab repository by following the link provided in the “Data Availability” section.

**Figure 7:**
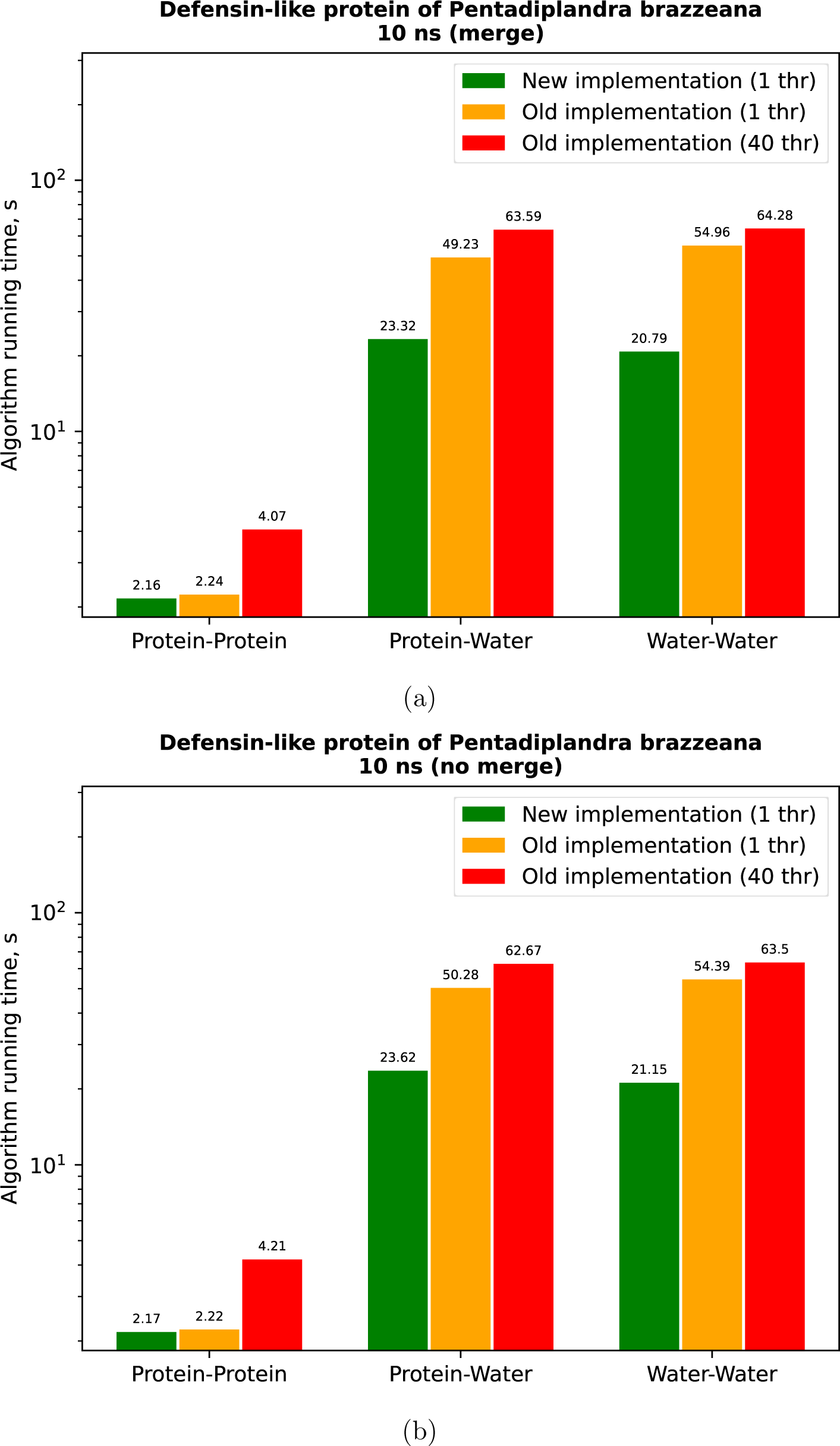
Comparison of the running time of the algorithms on the trajectory of the defensin-like protein Pentadiplandra brazzeana with a duration of 10 ns with (7a) and without (7b) the use of the hydrogen merge option.

**Figure 8:**
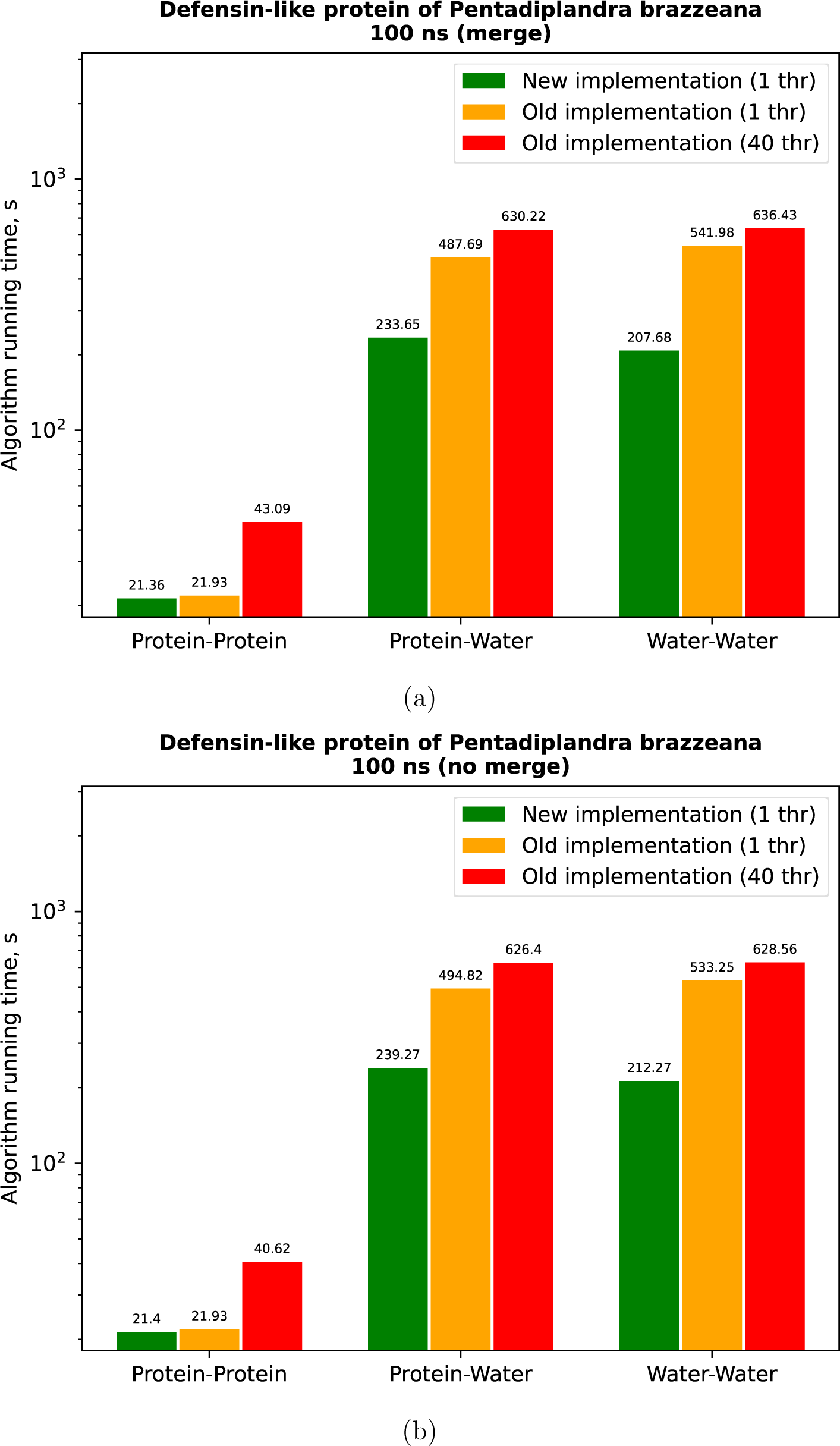
Comparison of the running time of the algorithms on the trajectory of the defensin-like protein Pentadiplandra brazzeana with a duration of 100 ns with (8a) and without (8b) the use of the hydrogen merge option.

**Figure 9:**
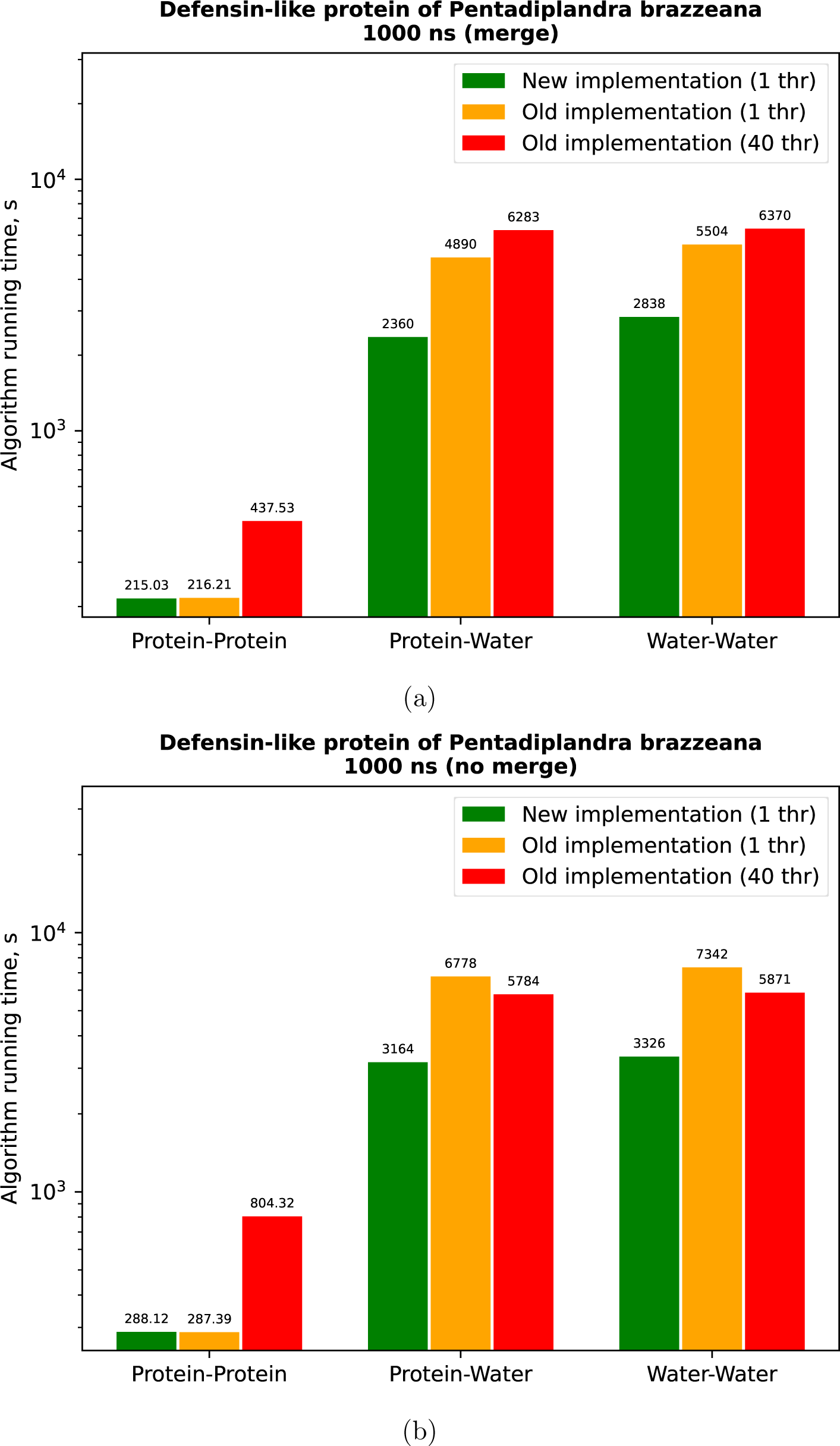
Comparison of the running time of the algorithms on the trajectory of the defensin-like protein Pentadiplandra brazzeana with a duration of 1000 ns with (9a) and without (9b) the use of the hydrogen merge option.

**Figure 10:**
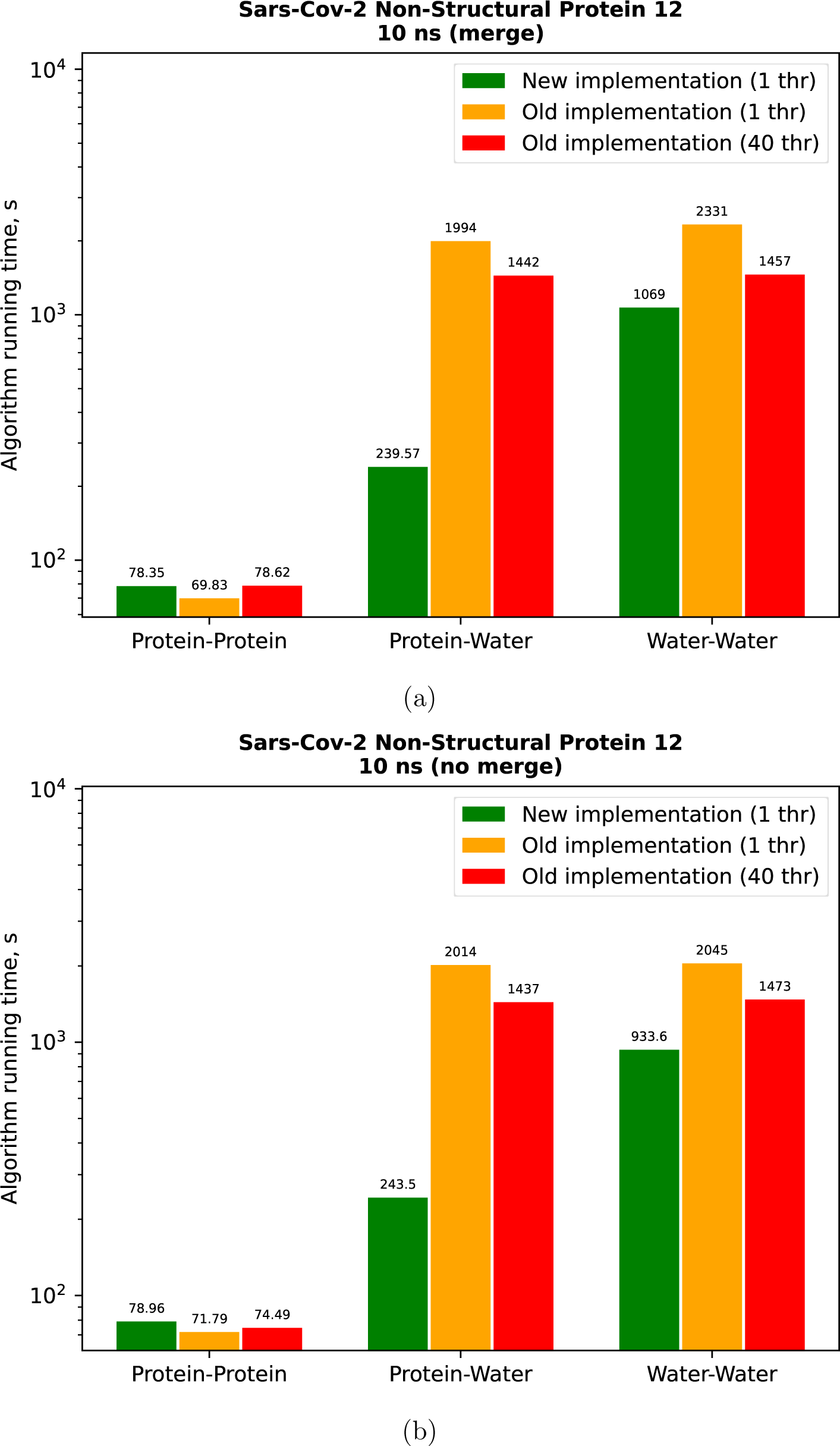
Comparison of the running time of the algorithms on the trajectory of Sars-Cov-2 Non-Structural Protein 12 with a duration of 10 ns with (10a) and without (10b) the use of the hydrogen merge option.

**Figure 11:**
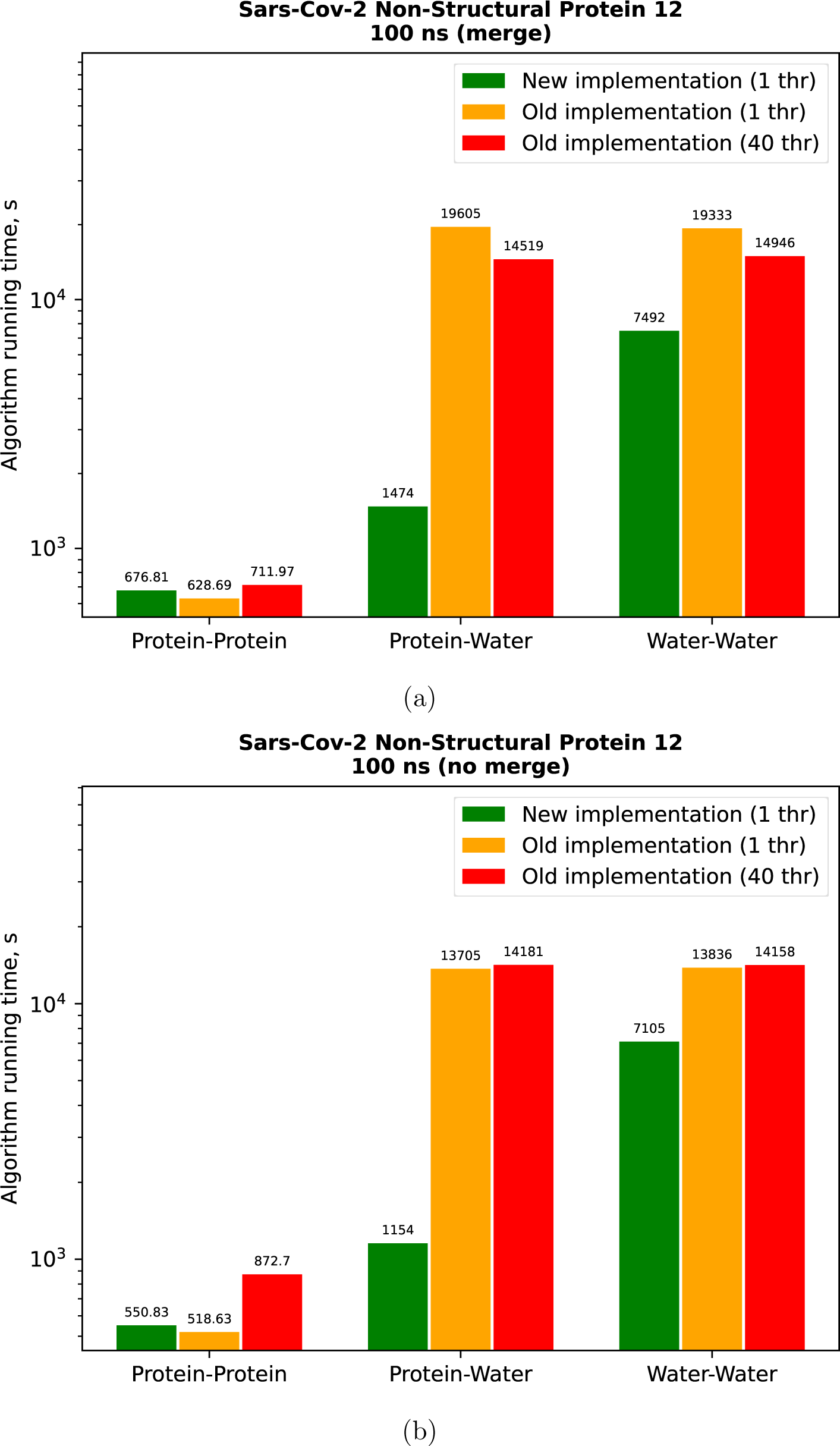
Comparison of the running time of the algorithms on the trajectory of Sars-Cov-2 Non-Structural Protein 12 with a duration of 100 ns with (11a) and without (11b) the use of the hydrogen merge option.

**Figure 12:**
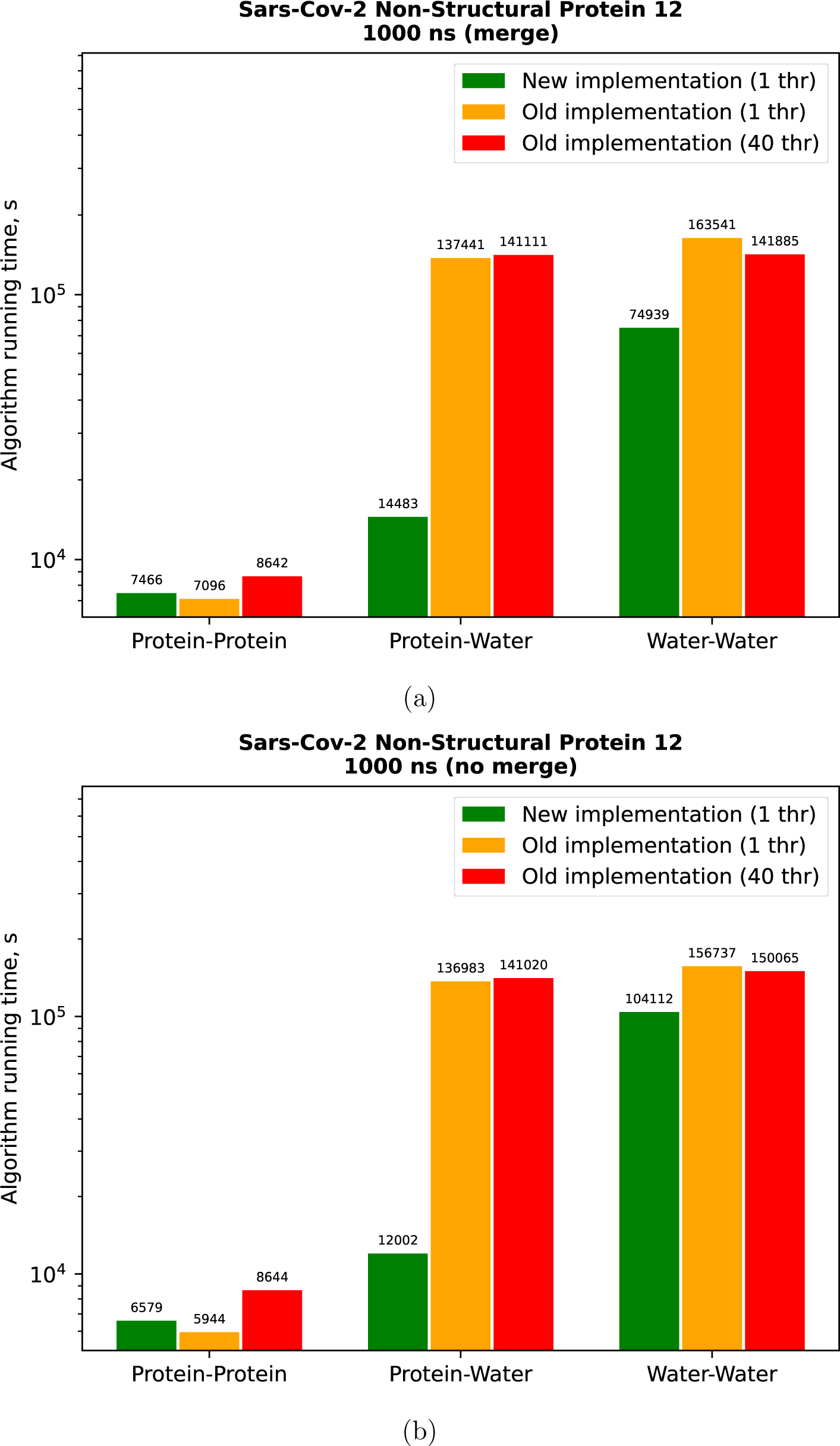
Comparison of the running time of the algorithms on the trajectory of Sars-Cov-2 Non-Structural Protein 12 with a duration of 1000 ns with (12a) and without (12b) the use of the hydrogen merge option.

**Figure 13:**
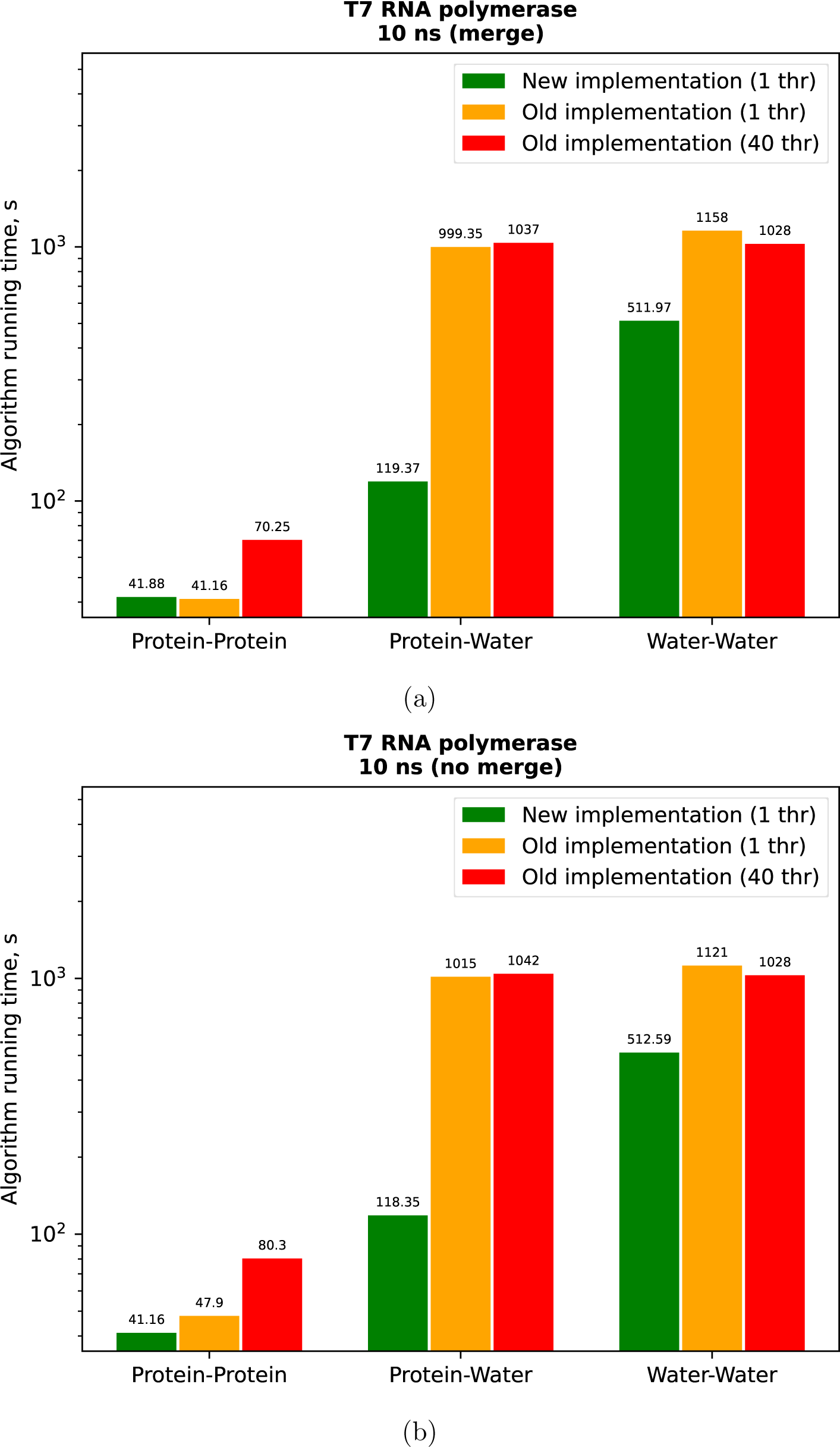
Comparison of the running time of the algorithms on the trajectory of T7 RNA polymerase with a duration of 10 ns with (13a) and without (13b) the use of the hydrogen merge option.

**Figure 14:**
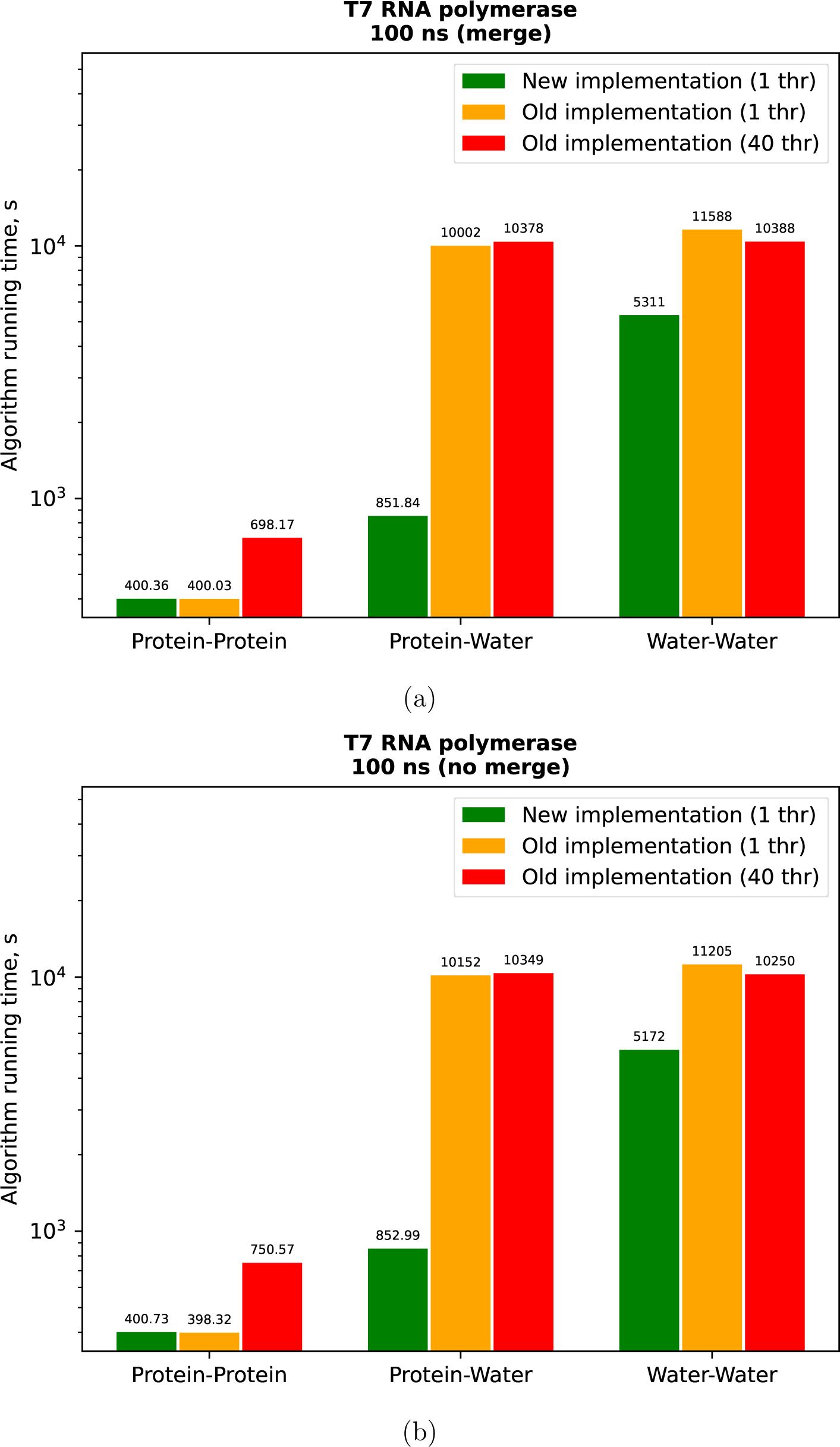
Comparison of the running time of the algorithms on the trajectory of T7 RNA polymerase with a duration of 100 ns with (14a) and without (14b) the use of the hydrogen merge option.

**Figure 15:**
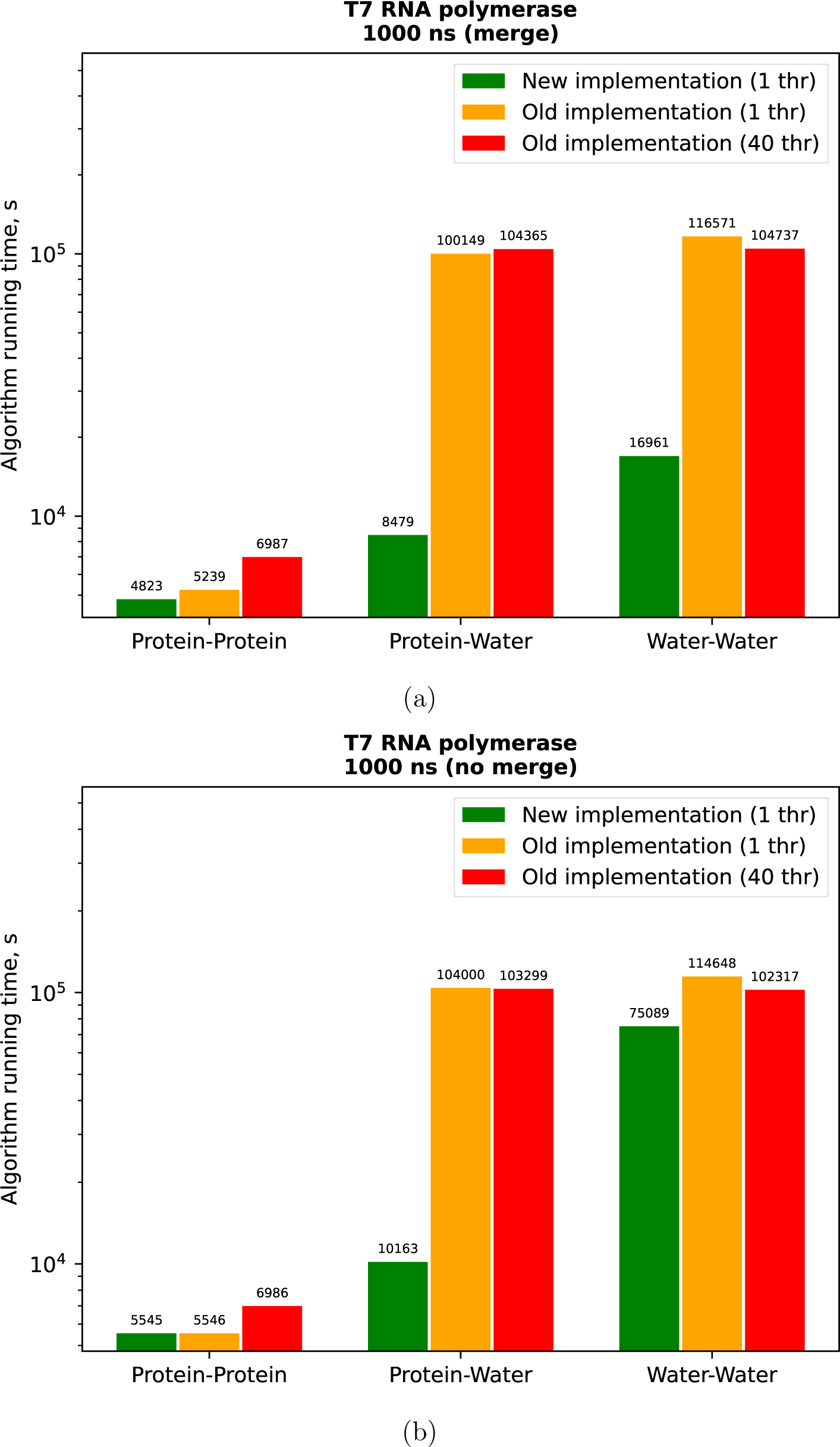
Comparison of the running time of the algorithms on the trajectory of T7 RNA polymerase with a duration of 1000 ns with (15a) and without (15b) the use of the hydrogen merge option.

**Figure 16:**
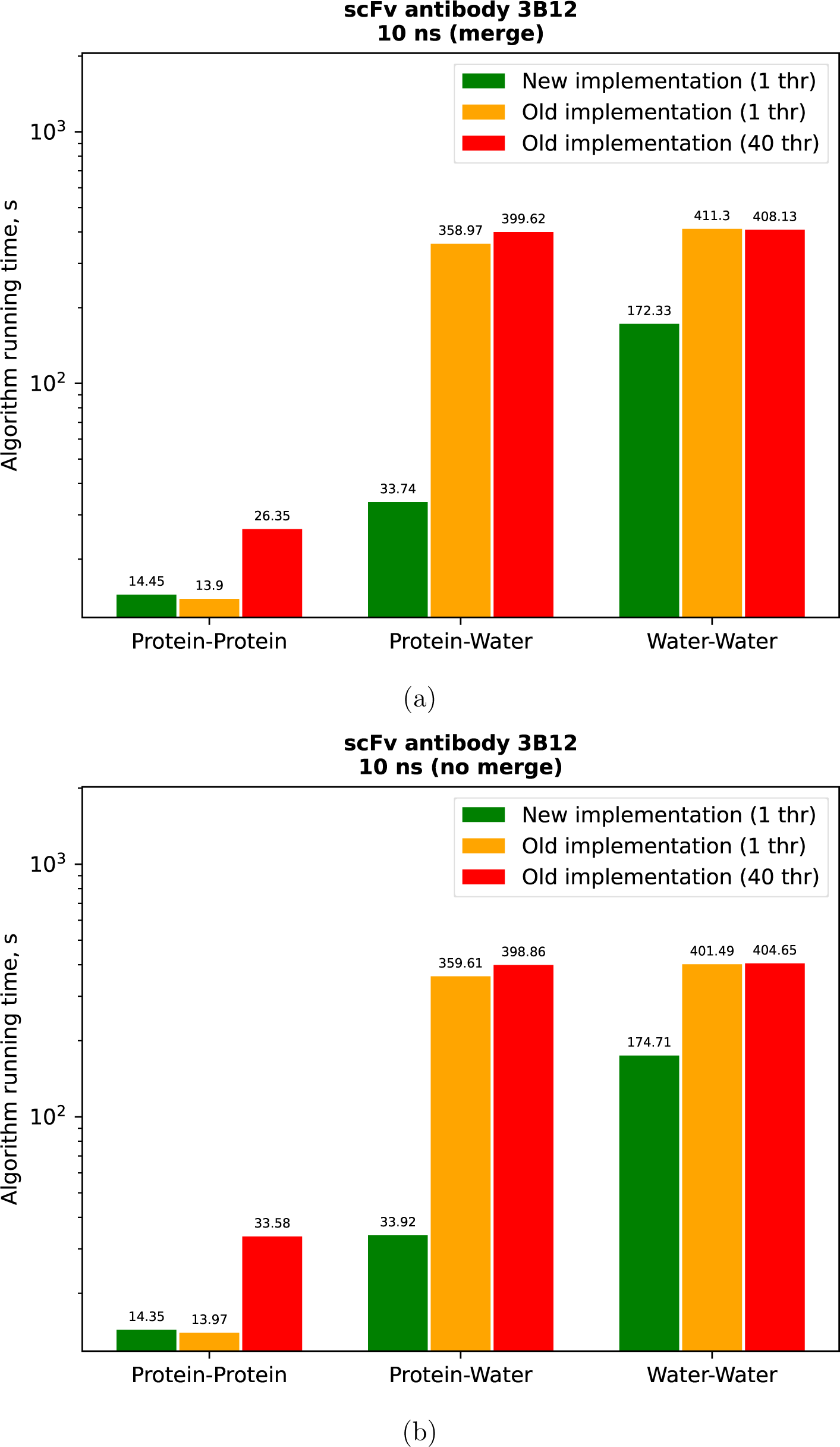
Comparison of the running time of the algorithms on the trajectory of scFv antibody 3B12 with a duration of 10 ns with (16a) and without (16b) the use of the hydrogen merge option.

**Figure 17:**
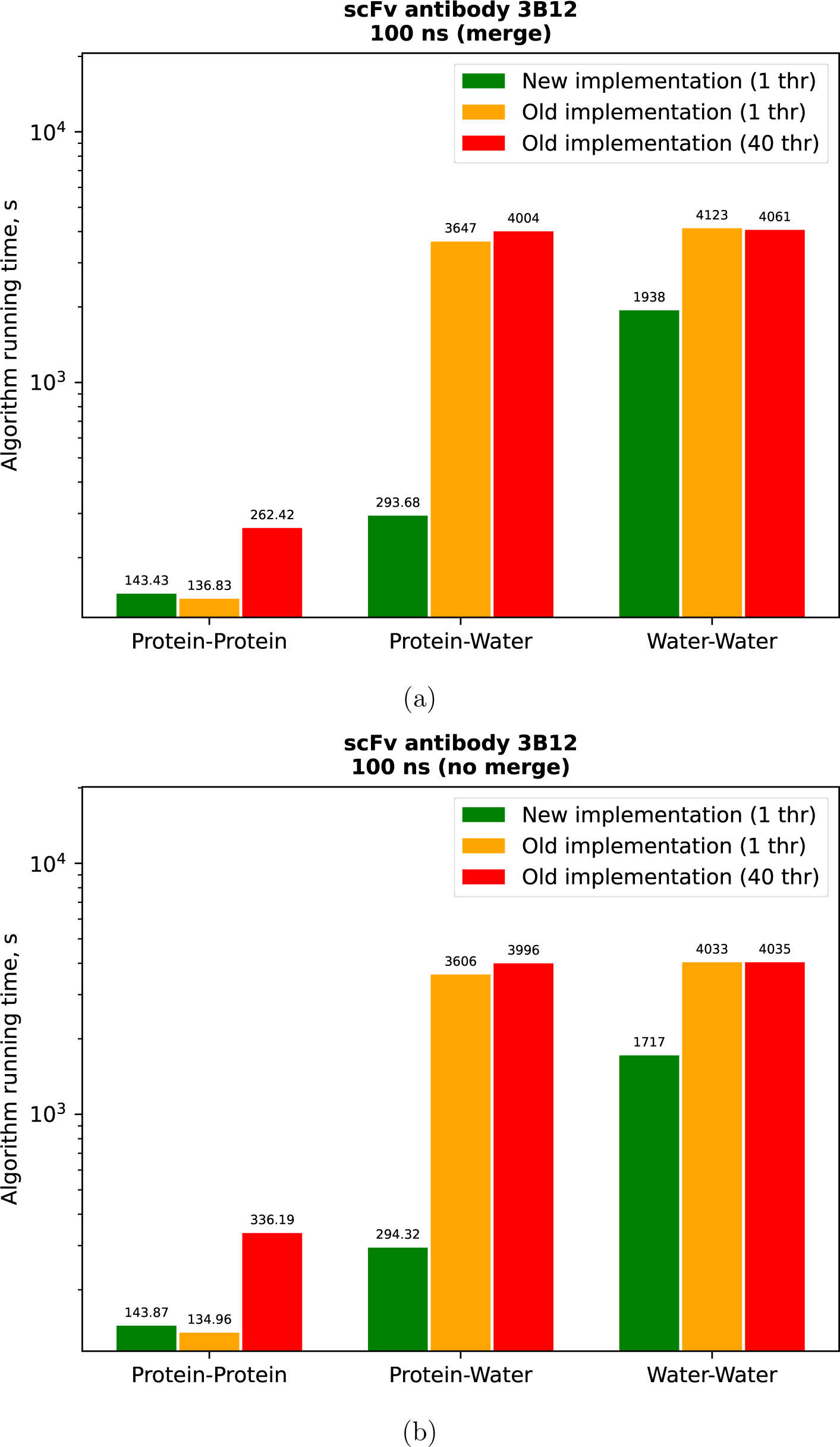
Comparison of the running time of the algorithms on the trajectory of scFv antibody 3B12 with a duration of 100 ns with (17a) and without (17b) the use of the hydrogen merge option.

**Figure 18:**
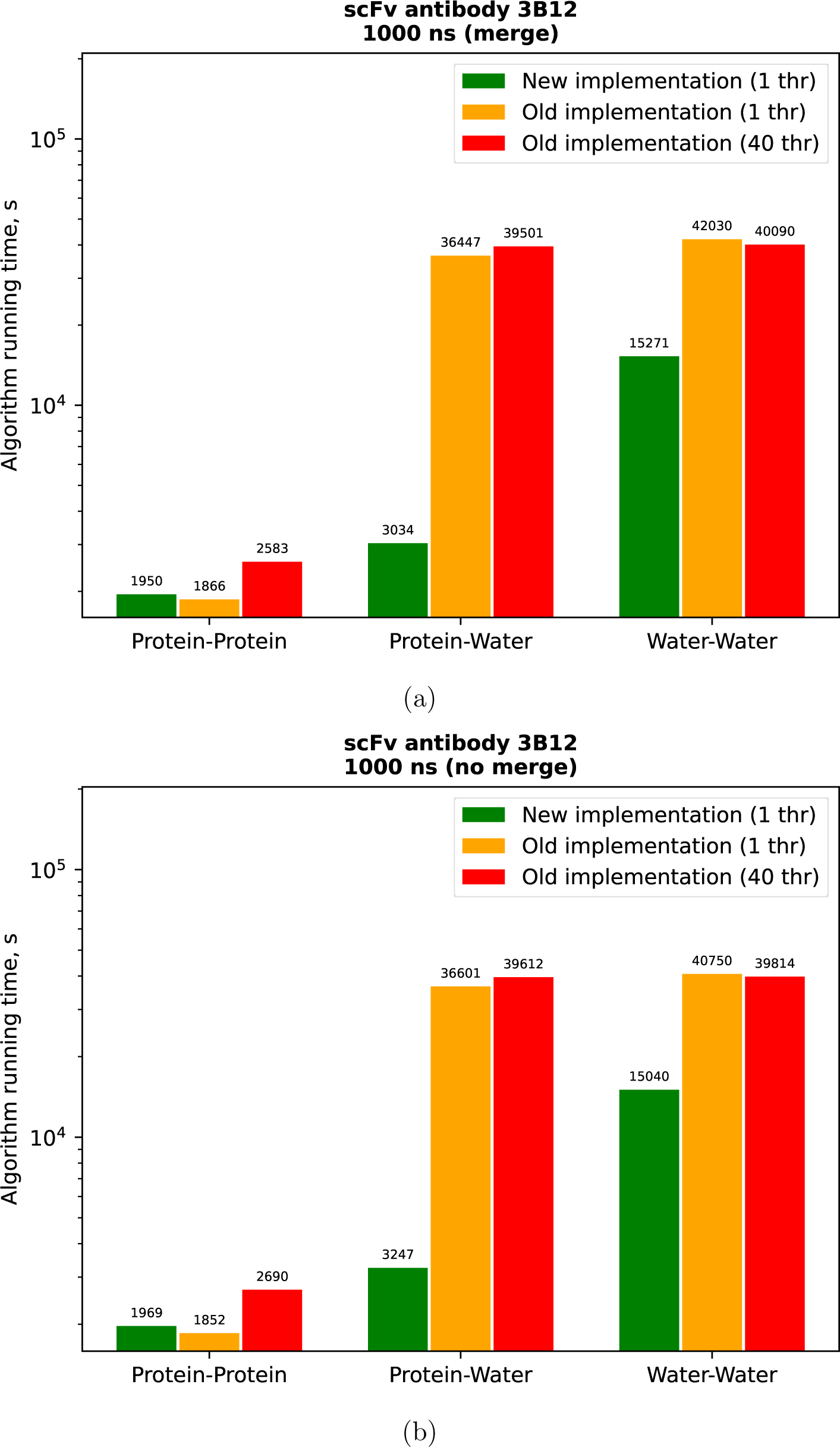
Comparison of the running time of the algorithms on the trajectory of scFv antibody 3B12 with a duration of 1000 ns with (18a) and without (18b) the use of the hydrogen merge option.

**Figure 19:**
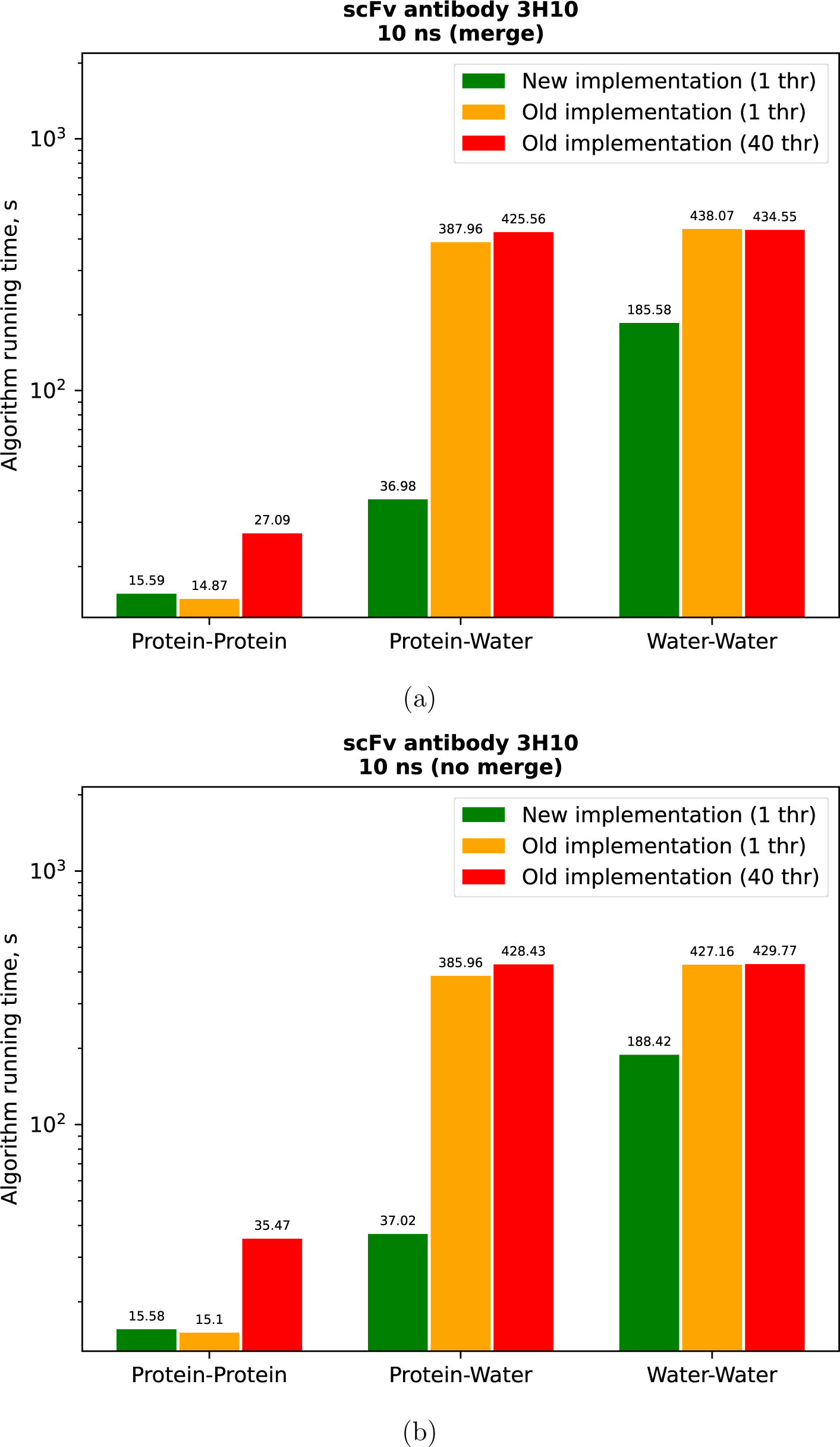
Comparison of the running time of the algorithms on the trajectory of scFv antibody 3H10 with a duration of 10 ns with (19a) and without (19b) the use of the hydrogen merge option.

**Figure 20:**
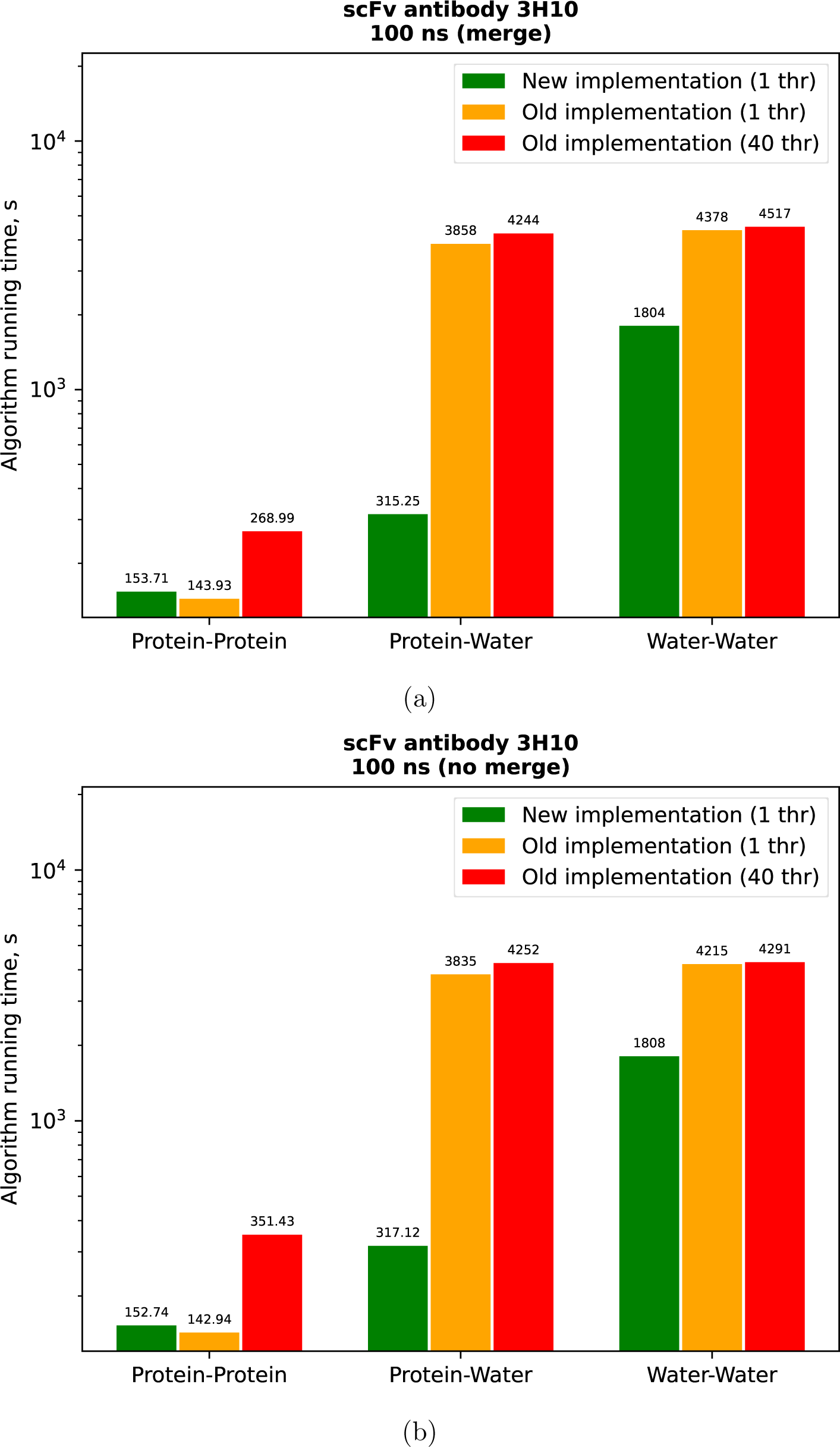
Comparison of the running time of the algorithms on the trajectory of scFv antibody 3H10 with a duration of 100 ns with (20a) and without (20b) the use of the hydrogen merge option.

**Figure 21:**
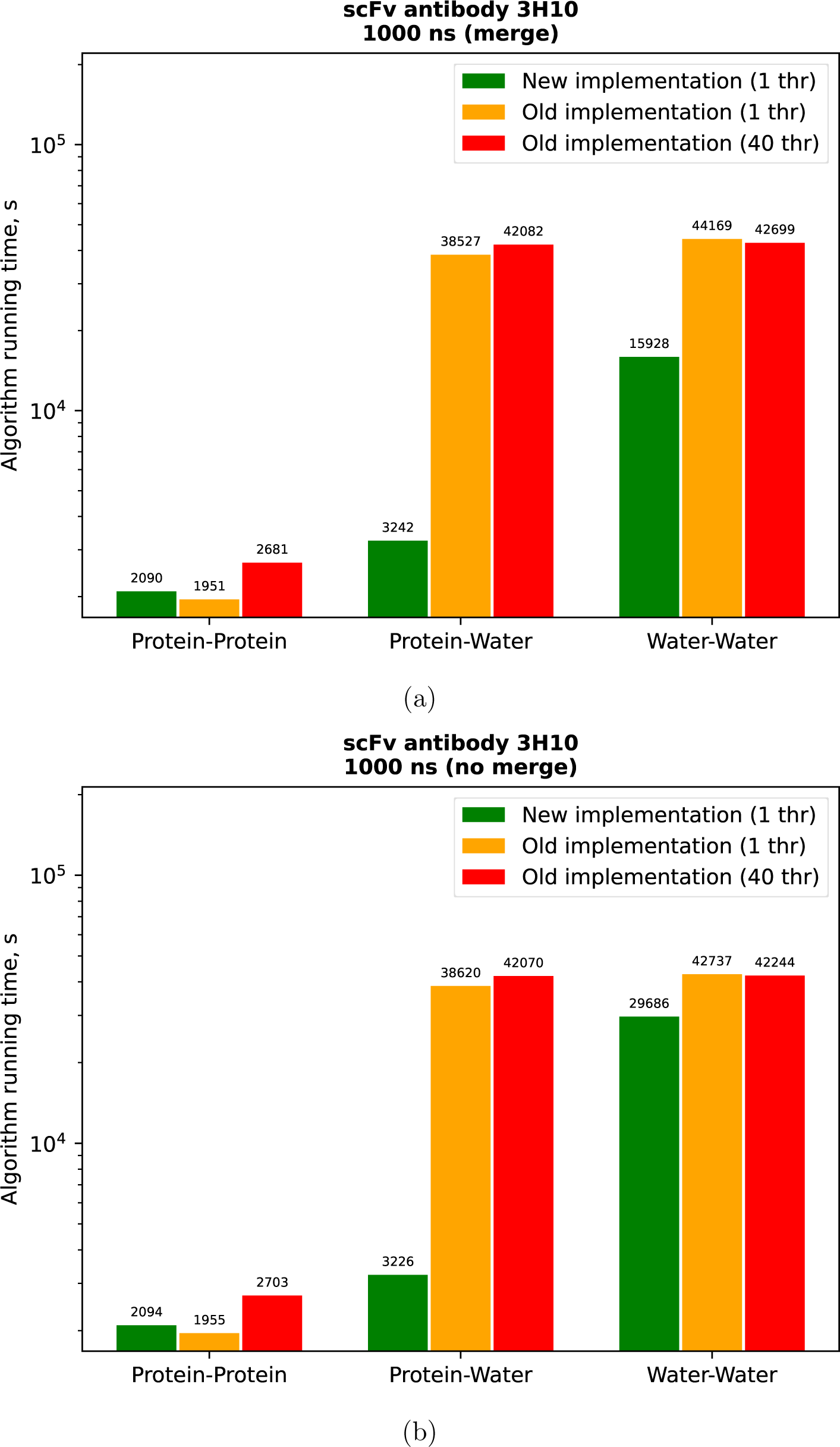
Comparison of the running time of the algorithms on the trajectory of scFv antibody 3H10 with a duration of 1000 ns with (21a) and without (21b) the use of the hydrogen merge option.

**Figure 22:**
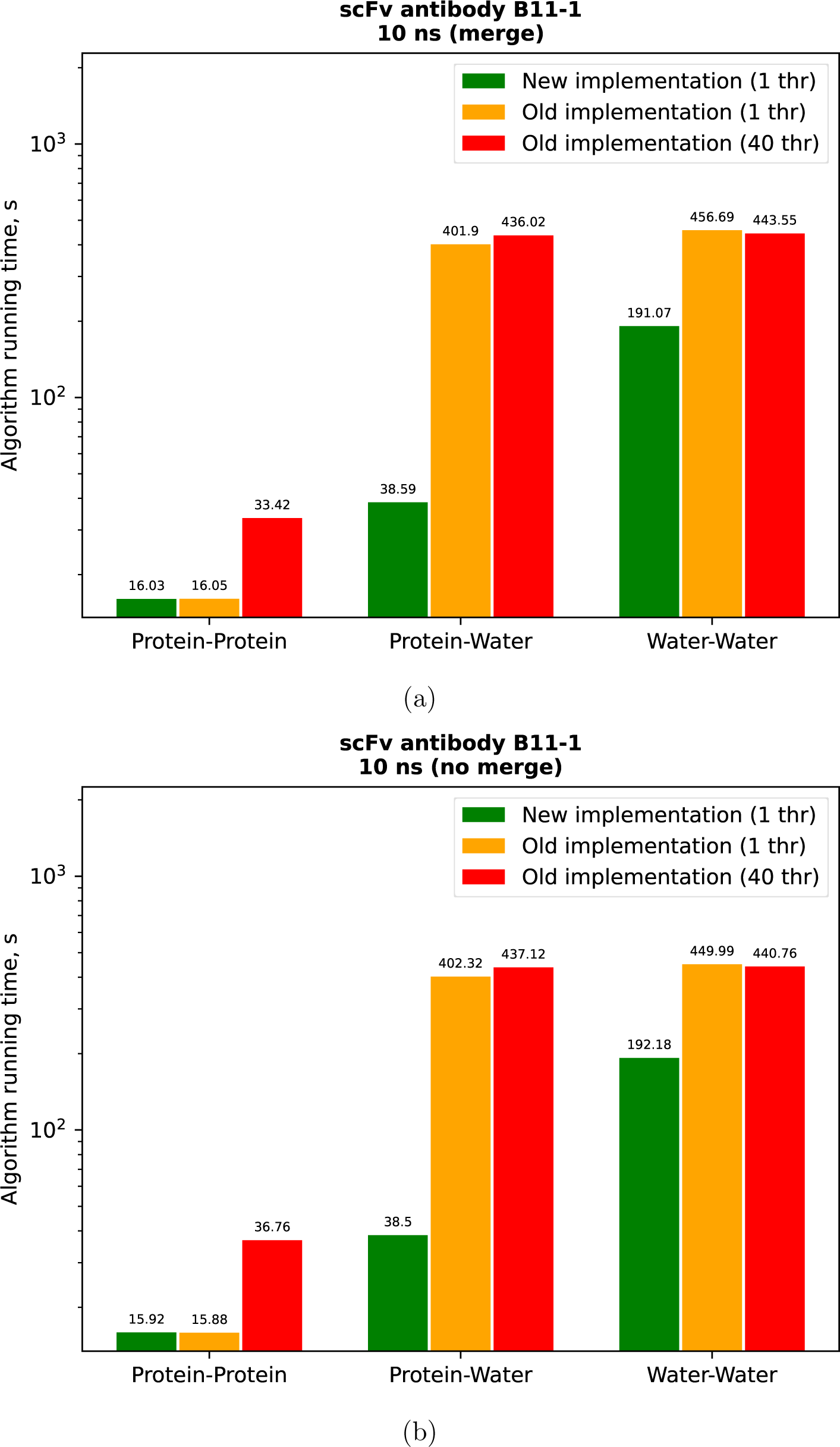
Comparison of the running time of the algorithms on the trajectory of scFv antibody B11-1 with a duration of 10 ns with (22a) and without (22b) the use of the hydrogen merge option.

**Figure 23:**
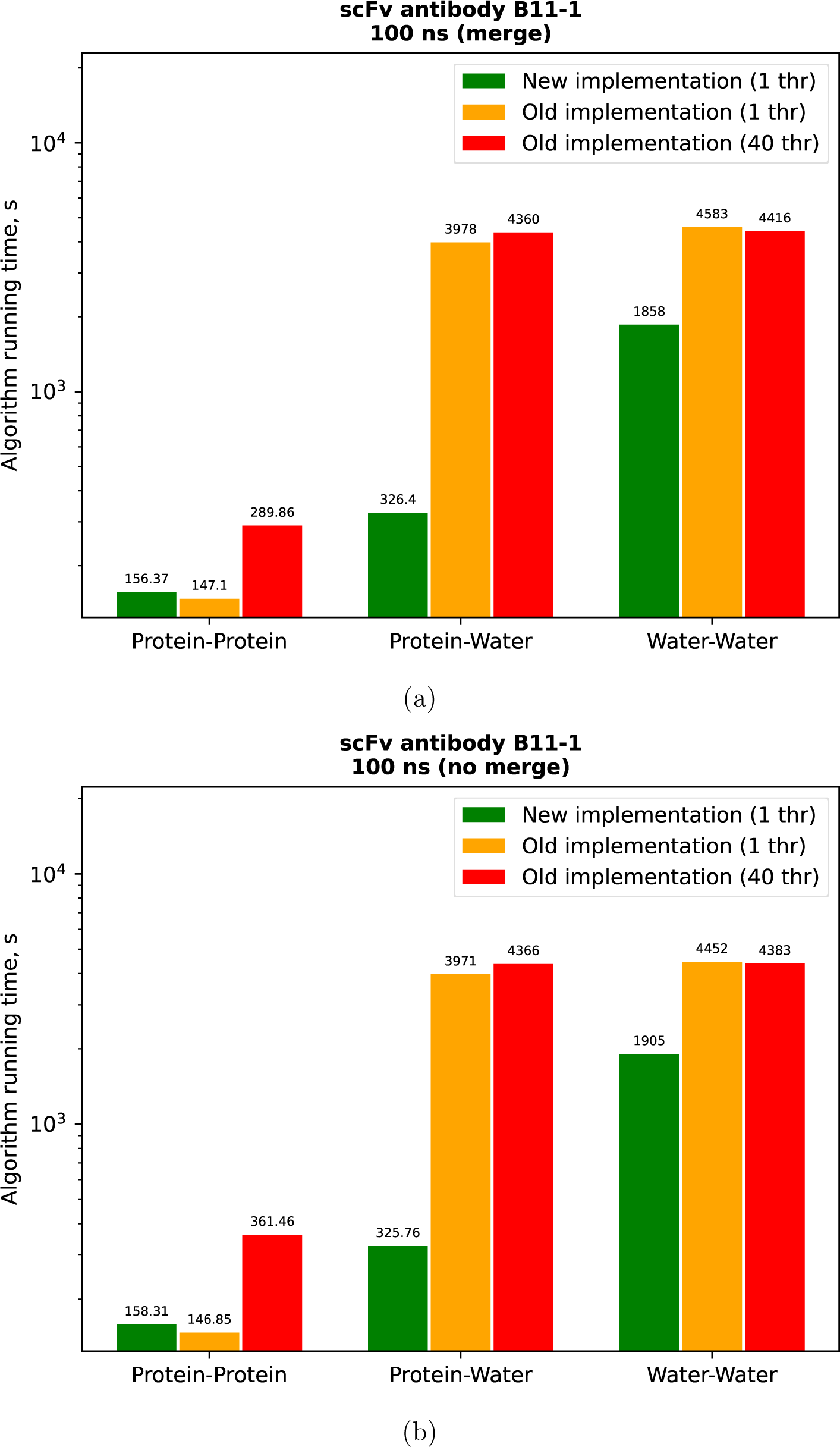
Comparison of the running time of the algorithms on the trajectory of scFv antibody B11-1 with a duration of 100 ns with (23a) and without (23b) the use of the hydrogen merge option.

**Figure 24:**
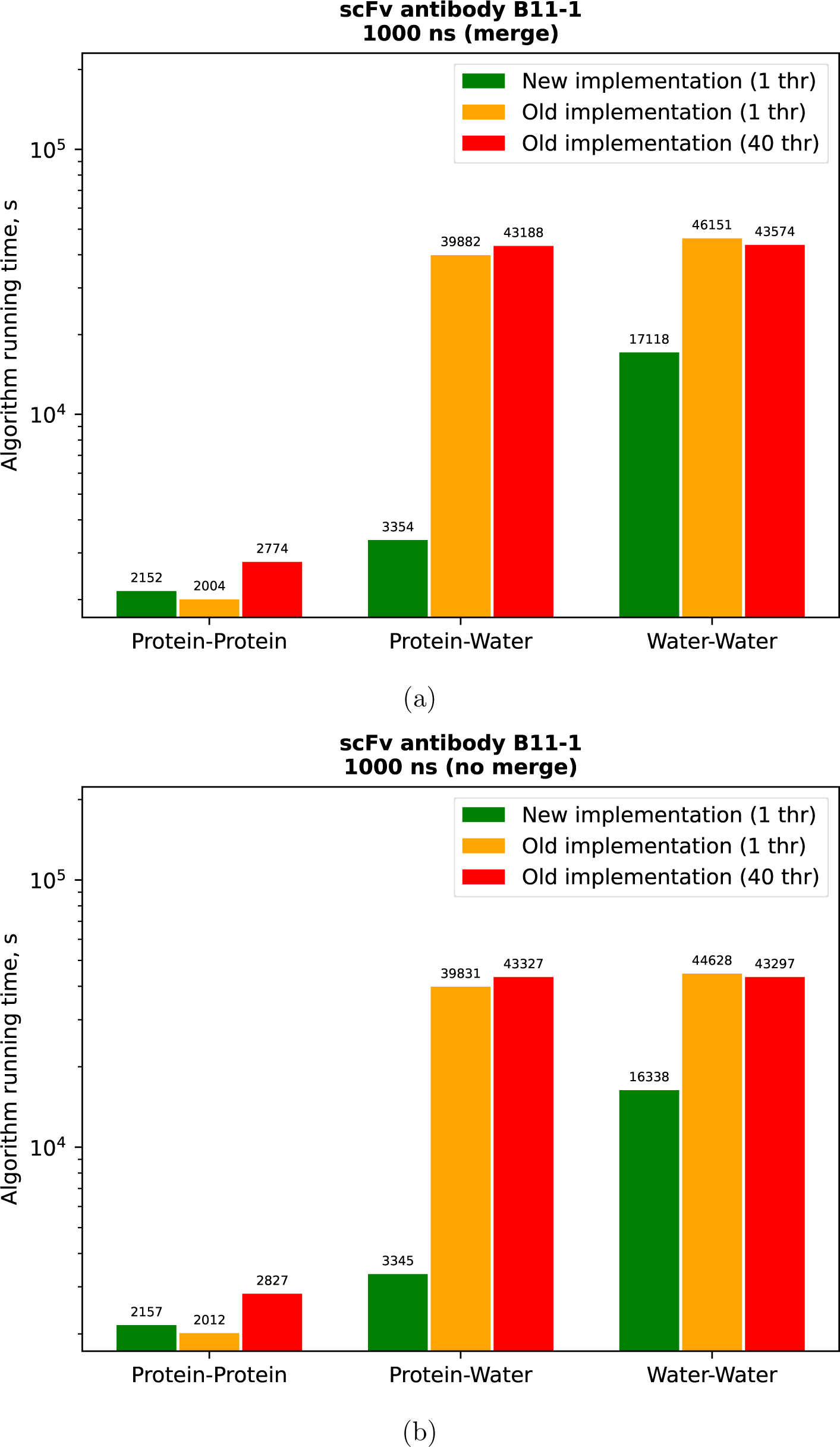
Comparison of the running time of the algorithms on the trajectory of scFv antibody B11-1 with a duration of 1000 ns with (24a) and without (24b) the use of the hydrogen merge option.

**Figure 25:**
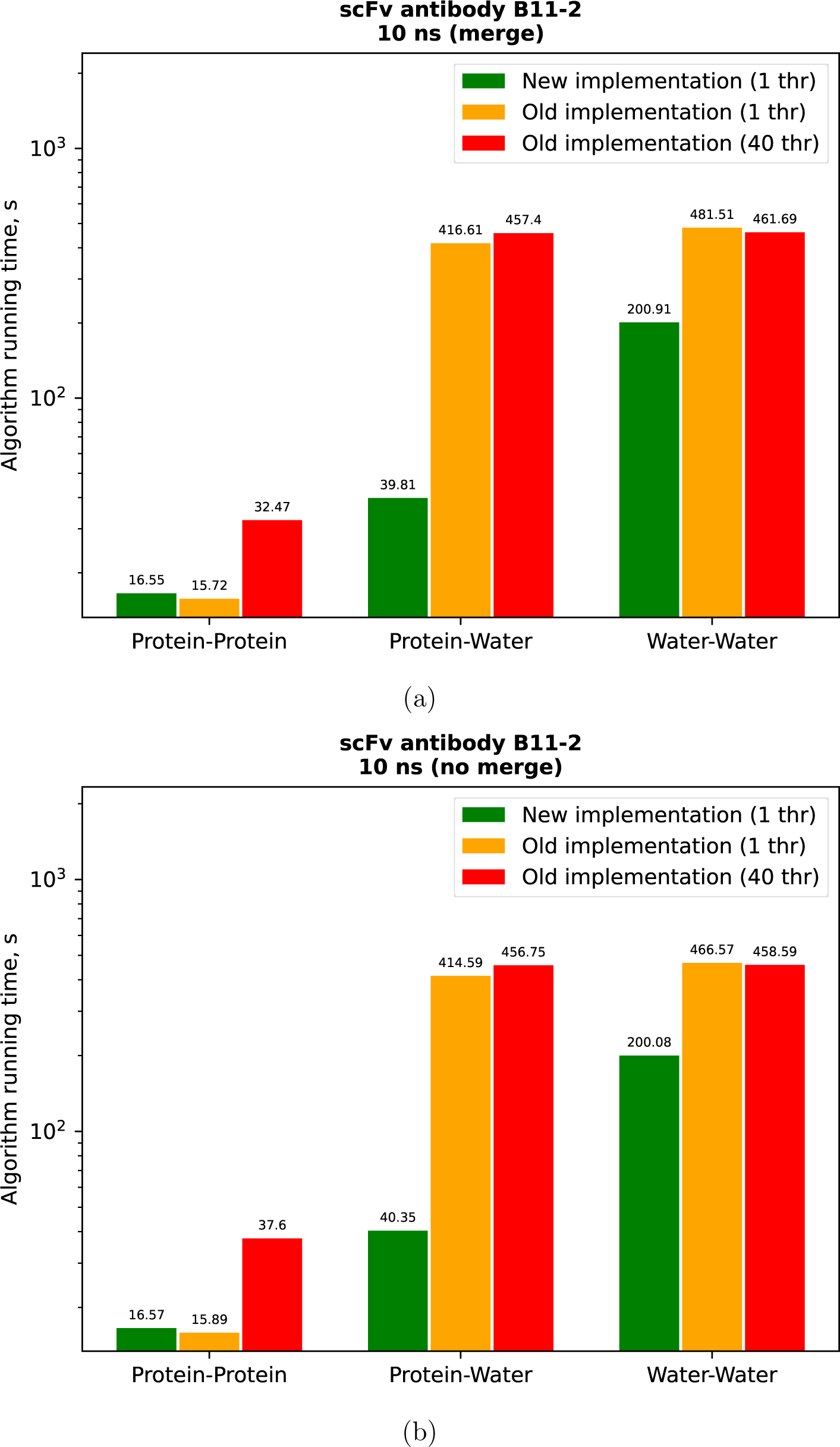
Comparison of the running time of the algorithms on the trajectory of scFv antibody B11-2 with a duration of 10 ns with (25a) and without (25b) the use of the hydrogen merge option.

**Figure 26:**
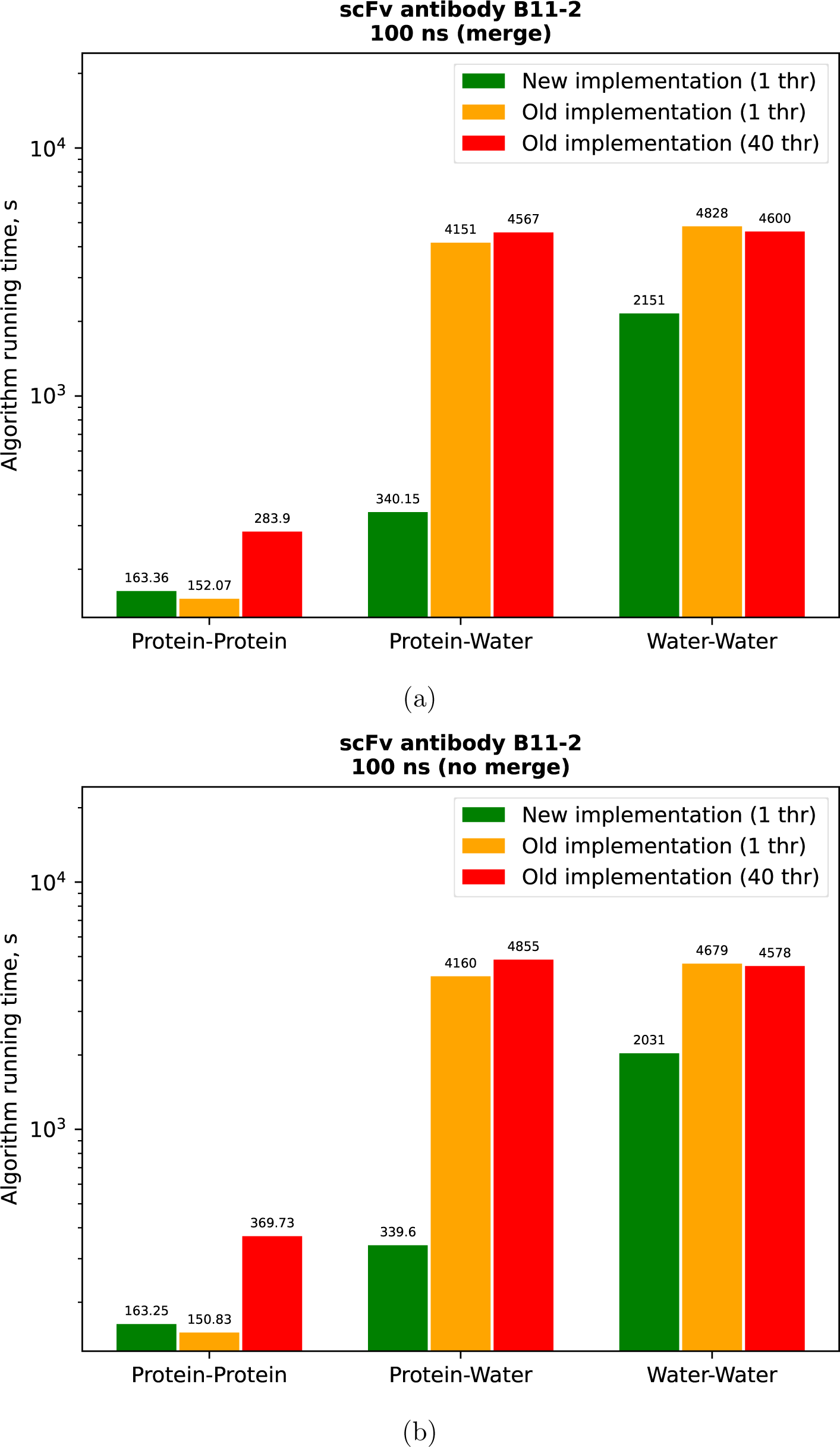
Comparison of the running time of the algorithms on the trajectory of scFv antibody B11-2 with a duration of 100 ns with (26a) and without (26b) the use of the hydrogen merge option.

**Figure 27:**
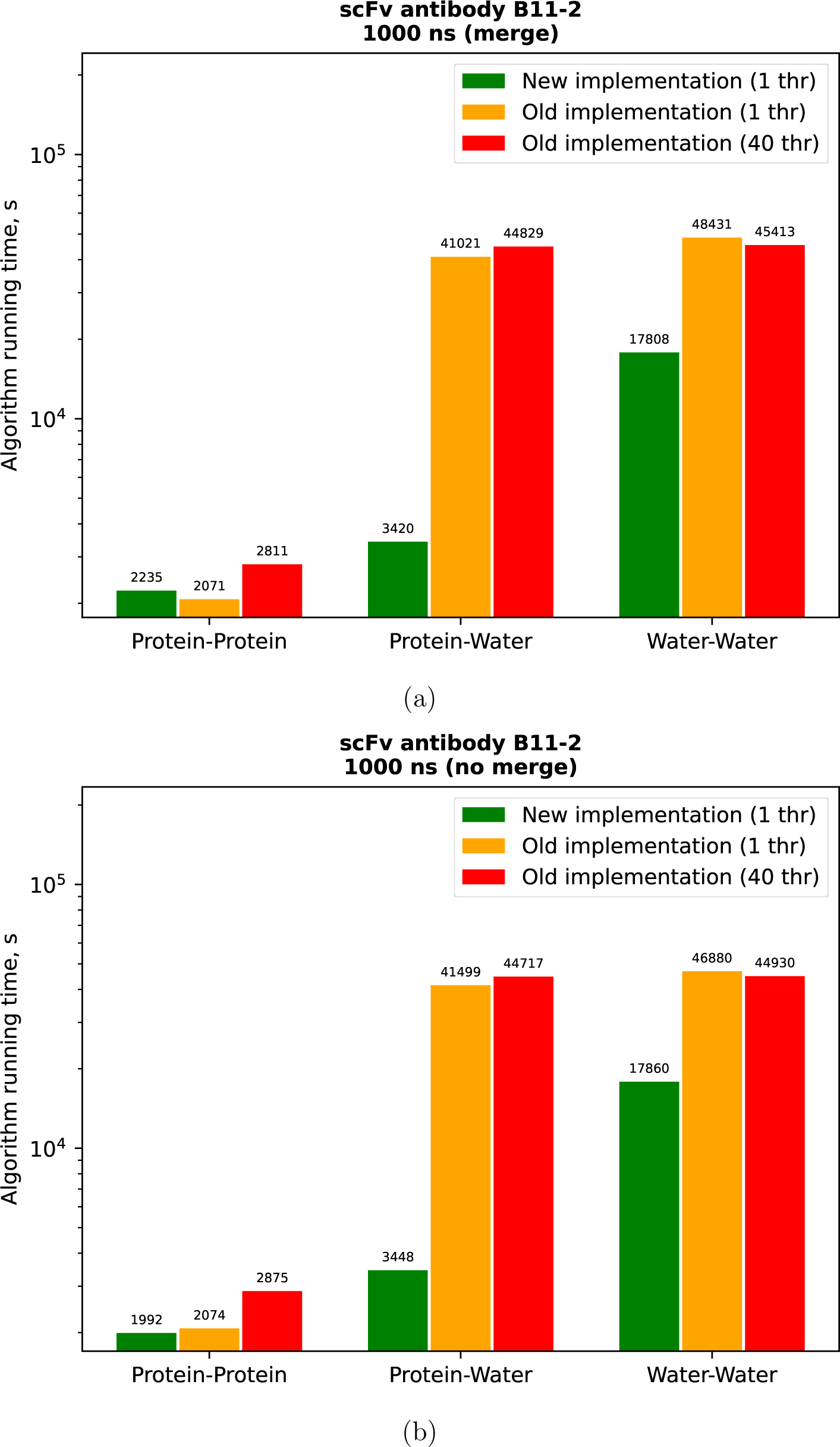
Comparison of the running time of the algorithms on the trajectory of scFv antibody B11-2 with a duration of 1000 ns with (27a) and without (27b) the use of the hydrogen merge option.

**Figure 28:**
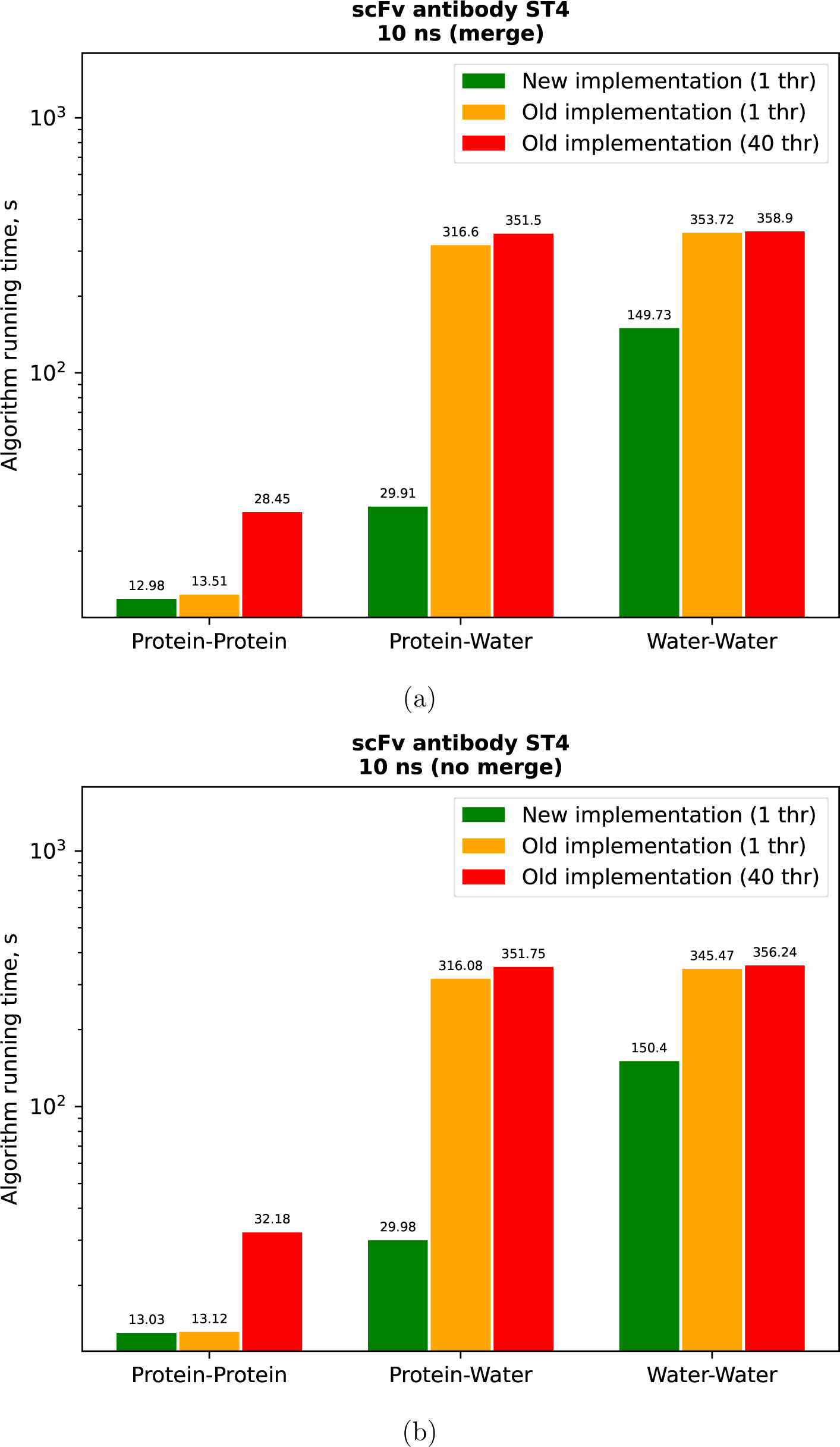
Comparison of the running time of the algorithms on the trajectory of scFv antibody ST4 with a duration of 10 ns with (28a) and without (28b) the use of the hydrogen merge option.

**Figure 29:**
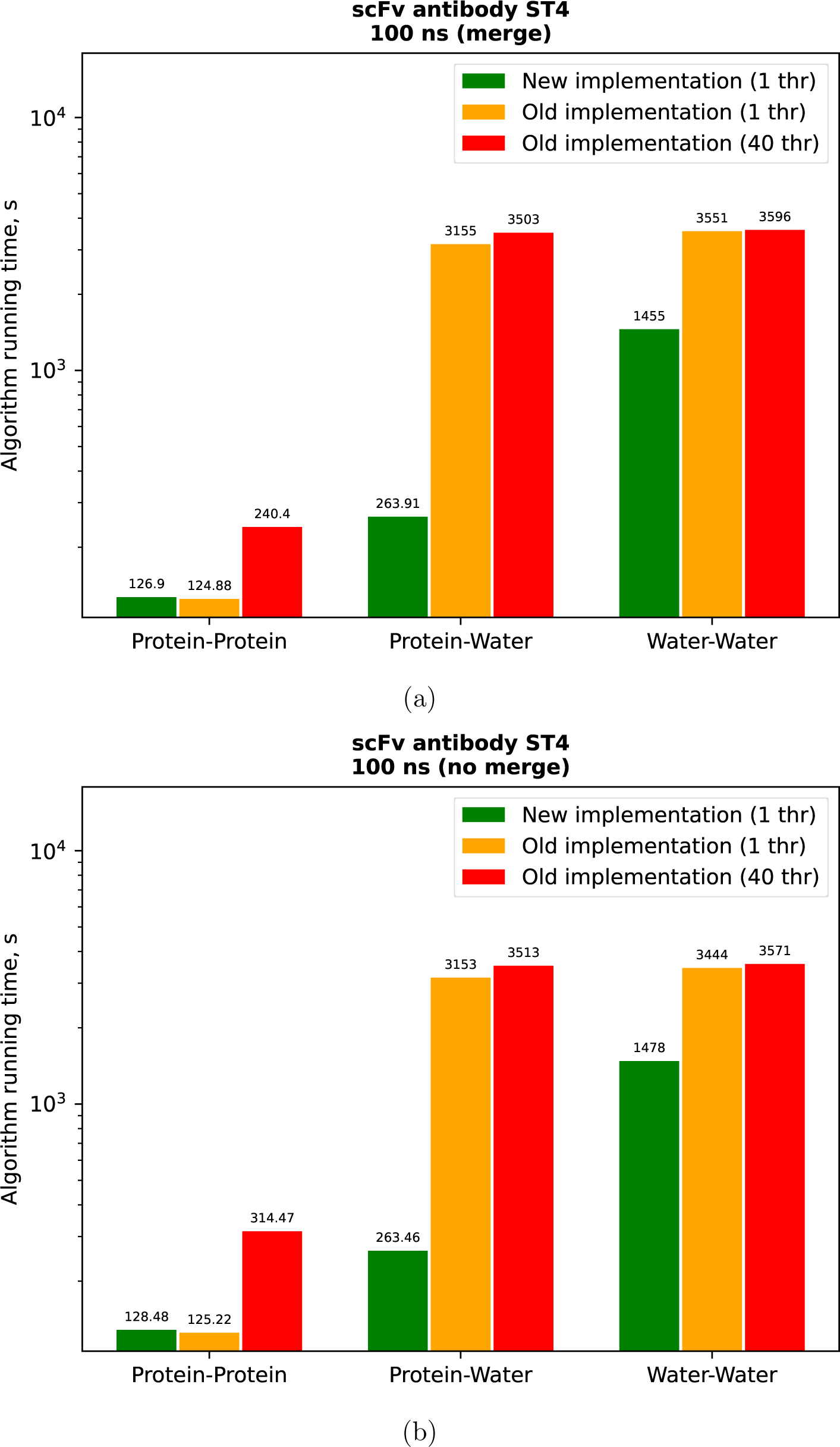
Comparison of the running time of the algorithms on the trajectory of scFv antibody ST4 with a duration of 100 ns with (29a) and without (29b) the use of the hydrogen merge option.

**Figure 30:**
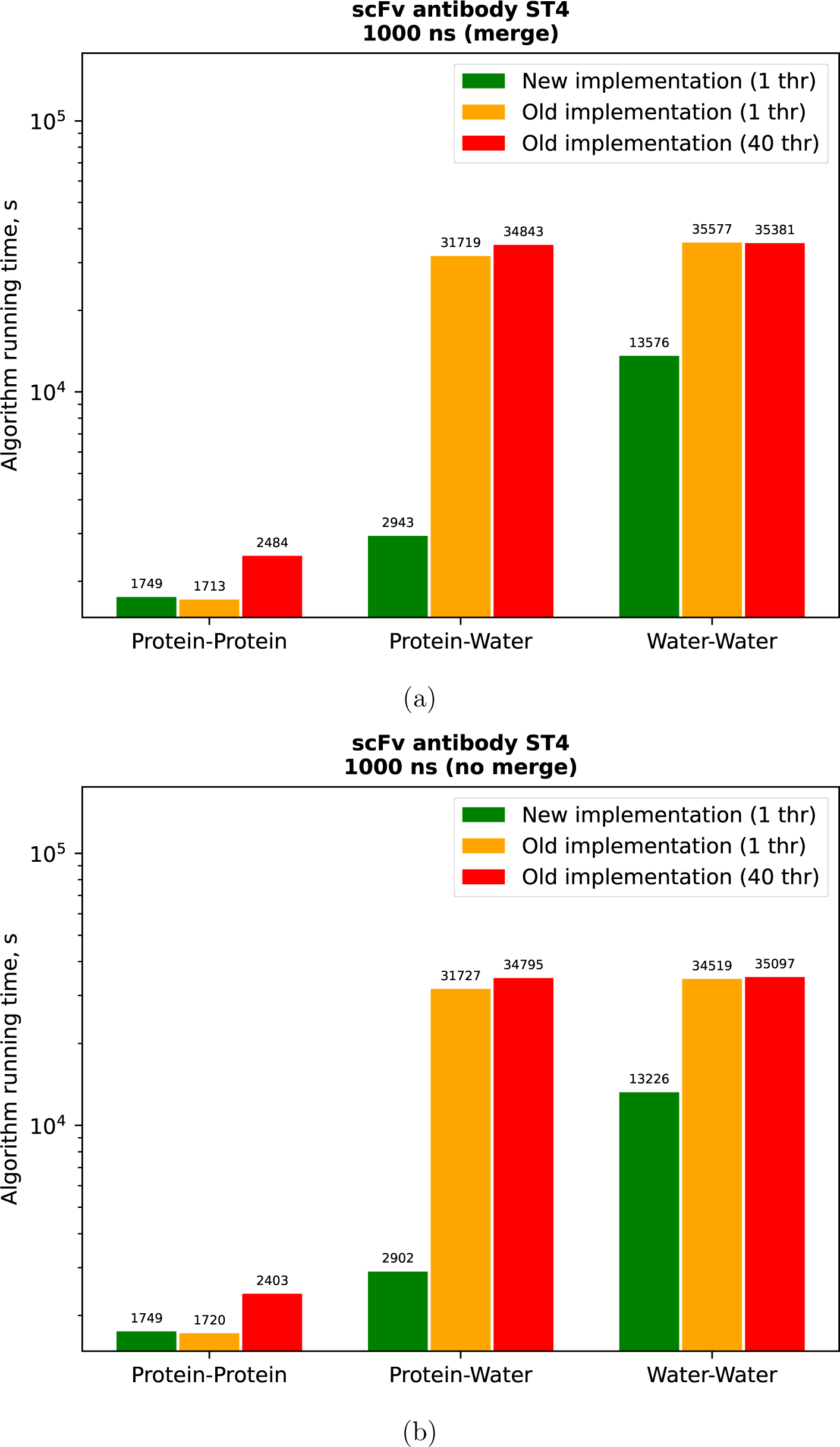
Comparison of the running time of the algorithms on the trajectory of scFv antibody ST4 with a duration of 1000 ns with (30a) and without (30b) the use of the hydrogen merge option.

Thus, according to the obtained data, we can conclude that the re-implementation of the algorithm for determining hydrogen bond networks works, in general, faster than the previous implementation. It is important to note that re-implementation is sometimes somewhat slower on small trajectories. This happens due to the peculiarities of the organization of the structures of the collections of atoms inside the trajectory analysis module. But, despite the slight delay on small trajectories, with an increase in the size of the trajectories arriving at the input, a significant difference in the speed of algorithm execution becomes noticeable in favor of the new implementation.

A formula for calculating the memory occupied for storing data on hydrogen bonds in the old and new implementations was derived. According to the calculations, the worst forecast for the amount of memory *V* (in bytes) occupied by the data store in the old implementation is:

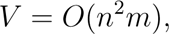

while in the new implementation it is:

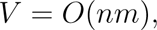

where *m* is the number of frames and *n* is the number of atoms in the investigated trajectory. From this we can conclude that the amount of occupied memory in the old implementation grows quadratically from the number of atoms, while in the new one it grows linearly. Therefore, in the old implementation the limiting factor is the total size of the system, while in the new it is the length of the trajectory. However, if in the old implementation this limitation cannot be bypassed, thereby becoming an insurmountable boundary for analysis, in the new implementation this limitation can be circumvented by splitting the trajectory under study into successive fragments. Additionally, the new implementation allows storing, in addition to the list of hydrogen bonds, the numbers of frames in which they were found, while in the old implementation this possibility was not implemented.

It’s also important to note that the old implementation supports multithreading, while the new implementation doesn’t (at the moment). However, it was noticed that when even 40 parallel threads were connected to perform calculations in large systems (such as the molecular dynamics trajectory for 10000 and 100000 frames), the running time of the old implementation was still significantly longer than the running time of the new implementation. It was noticed that with an increase in the number of threads in the old implementation, sometimes the calculation time increased when evaluating hydrogen bond networks within a protein and between a protein and water, although it was expected that an increase in the number of threads should, on the contrary, dramatically increase the calculation speed. This indicates the imperfection of the implementation of the algorithm in the old version. Despite all of the above, the creation of a multi-threaded mode for the new implementation is in the plans for the further development of scientific work in order to further reduce the calculation time of the developed software algorithm.

## References

(1) Helmenstine, A. M. Hydrogen Bond Definition and Examples. ThoughtCo 2023,

(2) Pace, C. et al. Contribution of hydrogen bonds to protein stability. Protein science : a publication of the Protein Society 2014, 23.

(3) Hubbard, R. E.; Haider, M. K. Hydrogen Bonds in Proteins: Role and Strength. 2010,

(4) Abraham, M. et al. GROMACS 2023.1 Manual (2023.1).

(5) Dannenberg, J. J. An Introduction to Hydrogen Bonding By George A. Jeffrey (University of Pittsburgh). Journal of the American Chemical Society 1998, 120, 5604–5604.

(6) Steiner, T. The Hydrogen Bond in the Solid State. Angewandte Chemie International Edition 2002, 41, 48–76.

(7) Yao, L.; Vögeli, B.; Ying, J.; Bax, A. NMR Determination of Amide N-H Equilibrium Bond Length from Concerted Dipolar Coupling Measurements. Journal of the American Chemical Society 2008, 130, 16518–16520.

(8) Demaison, J.; Herman, M.; Lievin, J. The equilibrium OH bond length. International Reviews in Physical Chemistry 2007, 26, 391–420.

(9) Jumper, J. et al. Highly accurate protein structure prediction with AlphaFold. Nature 2021 596:7873 2021, 596, 583–589.

(10) Abraham, M. et al. GROMACS 2023 Source code. 2023; 10.5281/zenodo.7588619.

(11) Maier, J. A.; Martinez, C.; Kasavajhala, K.; Wickstrom, L.; Hauser, K. E.; Simmerling, C. ff14SB: Improving the Accuracy of Protein Side Chain and Backbone Parameters from ff99SB. Journal of Chemical Theory and Computation 2015, 11, 3696–3713.

